# Lack of dominant-negative activity for tumor-associated ZNRF3 missense mutations at endogenous expression levels

**DOI:** 10.1101/2024.03.14.585013

**Authors:** Shanshan Li, Jiahui Niu, Ruyi Zhang, Sanne Massaar, Jenna van Merode, Nicky de Schipper, Lisa van de Kamp, Maikel P. Peppelenbosch, Ron Smits

**Affiliations:** Department of Gastroenterology and Hepatology, Erasmus MC Cancer Institute, University Medical Center Rotterdam, The Netherlands

**Keywords:** ZNRF3 and RNF43, Wnt/β-catenin signaling, neoplasms, mutation, missense, dominant-negative

## Abstract

ZNRF3, a negative regulator of β-catenin signaling, removes Wnt receptors from the membrane. Currently, it is unknown which tumor-associated variants can be considered driver mutations and through which mechanisms they contribute to cancer. Here we show that all truncating mutations analyzed at endogenous levels exhibit loss-of-function, with longer variants retaining partial activity. Regarding missense mutations, we show that 27/82 ZNRF3 variants in the RING and R-Spondin domain structures, lead to (partial) loss-of-function/hyperactivation. Mechanistically, defective R-spondin domain variants appear to undergo endoplasmic-reticulum-associated degradation due to protein misfolding. They show reduced stability and fail to reach the membrane correctly, which can be partially restored for several variants by culturing cells at 27°C. Although RING and R-spondin domain mutations in RNF43/ZNRF3 are often considered to possess dominant-negative oncogene-like activity in cancers, our findings challenge this notion. When representative variants are heterozygously introduced into endogenous ZNRF3, their impact on β-catenin signaling mirrors that of heterozygous knockout, suggesting that the supposed dominant-negative effect is non-existent. In other words, so-called “hyperactivating” ZNRF3/RNF43 mutations behave as classical loss-of-function mutations at endogenous levels. Taken together, our findings provide valuable information on ZNRF3 mutation impact in tumorigenesis and clarify their mechanism of action.

## Introduction

The Wnt/β-catenin signaling pathway is dysregulated in a large number of cancer types (1). Intracellularly, nuclear β-catenin signaling levels are regulated by a breakdown complex consisting of the APC tumor suppressor, scaffold proteins AXIN1/2 and the kinases GSK3β and CK1. Mutation of APC, AXIN1/2 or β-catenin itself, results in the formation of cancers with aberrantly increased β-catenin signaling, which have become largely independent of extracellular Wnt ligands (2). A second group of cancers acquires mutations in the homologous *RNF43* and *ZNRF3* genes. They encode for transmembrane E3-ubiquitin ligases that remove Wnt-receptors from the membrane, thereby limiting the level of Wnt-induced β-catenin signaling (2–5). An important feature of this subset of cancers is that increased β-catenin signaling is dependent on extracellular Wnt ligand exposure. This growth factor dependency is currently being exploited in clinical trials through the use of small molecule inhibitors against porcupine (PORCN), an O-acetyltransferase that is essential for Wnt-ligand secretion by cells (6, 7). Alternatively, treatments are aimed at preventing a direct interaction between Wnt ligands and their receptors (6, 7).

As these novel therapeutics progress into clinical trials, it is important to uncover which RNF43 and ZNRF3 variants observed in cancers, are likely to benefit from such treatments. For the *RNF43* gene this has already been studied more extensively. Truncating mutations that remove at least the C-terminal half of the protein, lead to a defective RNF43 protein, while longer truncated proteins retain partial functionality for which it is still unclear how much Wnt-dependency they confer onto cancers (8, 9). Missense RNF43 variants that affect β-catenin regulation are confined to the RING structure and extracellular R-spondin-binding domain (9). However, for basically all truncating and missense ZNRF3 variants their functional relevance is currently unknown, meaning that all tumor-associated ZNRF3 mutations are regarded as variants with uncertain significance (VUS).

Here, we aim to fill this knowledge gap by systematically profiling a large number of tumor-associated missense and truncating ZNRF3 variants. We find that at endogenous levels, all tested truncating mutations lose regulatory functions on Wnt/β-catenin signaling, with longer variants retaining some partial activity. Similar to RNF43, we identify various missense ZNRF3 mutations in the RING and R-spondin domains that lead to loss-of-function or hyperactivate β-catenin signaling. Previously, we and others suggested that this hyperactive behavior likely results from the formation of RNF43 and ZNRF3 homo- and heterodimers, in which the presence of one mutant RNF43/ZNRF3 inactivates the entire dimer complex in a dominant-negative manner (5, 9–12). In other words, hyperactivating RNF43 and ZNRF3 missense mutations may possess oncogene-like activity. However, this observation is entirely based on overexpression experiments, and for that reason we have tested this assumption here at endogenous expression levels. Furthermore, we show that missense variants in the R-spondin domain fail to reach the membrane correctly, a defect that could unexpectedly be partially rescued at low temperatures. Taken together, we identify a large number of ZNRF3 mutations that activate β-catenin signaling, and better define the mode of action through which ZNRF3 and RNF43 mutations contribute to cancer formation.

## Materials and methods

### Cell lines and culture

HEK293T cells and gene-edited variants thereof were maintained in DMEM culture medium supplemented with 10% fetal bovine serum (FBS) at 37°C in a humidified atmosphere containing 5% CO2. The culture medium was changed every 2-3 days. Mycoplasma tests showed that the cell lines were mycoplasma negative based on the real-time PCR method at Eurofins GATC-Biotech (Konstanz, Germany). L-Wnt3a and L-control conditioned medium was prepared according to standard procedures. The generation of RNF43-KO, ZNRF3-KO and double knockout HEK293T clones has been described previously (8). Identity of all cell lines and clones thereof, was confirmed by the Erasmus Molecular Diagnostics Department, using Powerplex-16 STR genotyping (Promega, Leiden, The Netherlands).

### Construction of ZNRF3 and RNF43 variant expression plasmids

Expression vectors for C-terminal FLAG-tagged ZNRF3-short and ZNRF3-long isoforms were generated using the C-terminal FLAG-tagged RNF43 as basis (8). The full-length cDNA of the human ZNRF3 was acquired from the human CRC DLD-1 cell line by RNA purification using NucleoSpin RNA II kit (Macherey Nagel) and reverse transcription RT-PCR using specific primers and Primescript RT reagent kit (TaKaRa). Then the ORF of the ZNRF3 short variant replaced RNF43 and was assembled into the C-terminal FLAG-tagged vector using Q5 Hot Start High-Fidelity DNA Polymerase (New England Biolabs) and the GeneArt Gibson Assembly HiFi Master Mix (Life Technologies) according to the manufacturer’s protocol. The long ZNRF3 variant was generated by Gibson cloning the 300 bp ORF of exon 1 into the short ZNRF3 expression plasmid. The 3xFLAG- and HA-tagged ZNRF3 expression vectors were generated with the Q5 Hot Start High-Fidelity DNA Polymerase Kit (New England Biolabs) by replacing the FLAG-tag. Both tags are preceded by a SGGGSGGGSG linker sequence.

To introduce missense and nonsense variants in the long ZNRF3 expression plasmid, two approaches were used. The first one made use of Q5 mutagenesis following the manufacturer’s instructions (New England Biolabs), but due to the high GC-content of exon1 specific for the ZNRF3 long isoform, all PCRs failed and variants were first introduced in the short ZNRF3 plasmid. Next, by using conventional restriction enzyme digests and ligation, exon1 was reintroduced. As a second system we made use of the Telesis BioXp system (TelesisBio, San Diego, CA) to directly synthesize and clone 32 additional R-spondin and 8 RING domain variants. These were cloned into NotI/BamHI-digested ZNRF3-HA and XbaI/BstEII-digested ZNRF3-3xFLAG plasmids, respectively.

Various RNF43 expression plasmids had been generated previously (8, 9, 13). In addition, 28 novel RNF43 RING domain variants were generated using Q5 mutagenesis (New England Biolabs). All ZNRF3 and RNF43 expression plasmids were full-length sequence verified. See **Supplementary Table 2 - 4** for primer sequences.

### Quantitative PCR

Total RNA was extracted using a NucleoSpin RNA kit (Macherey Nagel) and reverse transcribed with the Primescript RT reagent kit (TaKaRa) according to the manufacturer’s instructions. Quantitative PCR was performed in StepOne Real-Time PCR System (Applied Biosystems). All experiments were performed in triplicates. Gene expression analysis was performed using the comparative ΔΔC_T_ method with the housekeeping gene *GAPDH* for normalization. Primer sequences are provided in **Supplementary Table 5**.

### Luciferase reporter assays

Wnt Responsive Element (WRE) luciferase assays were executed using approximately 20% confluent HEK293T cells in 24-well plates. ZNRF3 variant plasmids (100ng), WRE plasmid (100ng), and CMV-Renilla (10ng) were transfected using Lipofectamine 2000 (Life Technologies). After transient transfection for 6 hours, cells were stimulated with L-Wnt3a or L-Control conditioned medium, and cultured for an additional two days. Firefly and Renilla luciferase activity was measured with the Dual-Luciferase Reporter Assay system (Promega) according to the manufacturer’s instruction in a LumiStar Optima luminescence counter (BMG LabTech, Offenburg, Germany). Firefly luminescence was normalized to Renilla. Transfections were performed in triplicate.

### Cell surface biotinylation and immunoprecipitation

HEK293T cells in six-well plates were transiently transfected using Lipofectamine 2000 at approximately 20% confluency with 600ng of C-terminal 3xFLAG-tagged RNF43 or ZNRF3 plasmids and cultured for 48h. Cells were washed with cold PBS once, and surface proteins were biotinylated with 0.5 mg/ml Premium Grade Sulfo-NHS-LC-Biotin (Fisher Scientific) in PBS at 4°C for 30 min. PBS with 100mM glycine was used to wash the cells once to quench and remove excess Biotin reagent and by-products. Then cells were washed twice with cold PBS and lysed with 500ul of IP buffer (30 mM Tris-HCl, 150 mM NaCl, 5 mM EDTA, 5 mM NaF, and 1% TritonX-100) containing Halt Protease and Phosphatase Inhibitor Single-use Cocktail (Thermo Fisher Scientific) and incubated 15 min on ice. Cell debris was collected by scraping and transferred to low adhesion tubes, followed by centrifugation at 11,000 g for 15 min at 4 °C. As input control, 10% of the cleared lysate was taken and mixed with 2xLaemmli with 0.1 mol/L Dithiothreitol (DTT), followed by heating for 7 minutes at 95 °C. The remaining supernatant was incubated with 50 μl of precleared anti-FLAG M2 Affinity Agarose Gel (cat. # A2220, Sigma-Aldrich) for 2 hours at 4 °C with rotating. After five times extensive washing with IP buffer, the immunoprecipitated bead conjugates were heated in 2xLaemmli/DTT sample buffer (95 °C, 7 min) and subjected to SDS-PAGE gel, followed by immunoblotting.

### Western blots

For western blot experiments, approximately 20% confluent HEK293T cells seeded in 12-wells plates were transfected with 300ng ZNRF3 encoding plasmid and 50ng GFP expression plasmid (pEGFP-C1, Clontech) using Lipofectamine 2000. Two days after transfection, cells were washed twice with PBS and lysed in 150μl 2xLaemmli/DTT buffer. Subsequently, collected samples were boiled for 10 minutes at 95°C. For immunoblotting, membranes were blocked with Odyssey blocking buffer (LI-COR-Bioscience, Lincoln, NE, USA). Mouse anti-FLAG (Anti-FLAG M2 mouse monoclonal antibody, F1804-1 Sigma-Aldrich, 1:1000), mouse anti-HA (Monoclonal Anti-HA antibody produced in mouse, H9658 Sigma-Aldrich, 1:1000) and rabbit anti-GFP (polyclonal anti-GFP rabbit IgG, A11122 Invitrogen, 1:000, or polyclonal anti-GFP rabbit IgG, A-6455 ThermoFisher, 1:2000) antibodies were then used. Biotin was detected using IRDye 800CW Streptavidin (1:1000, cat.#926-32230, Licor-Biosciences). Secondary antibodies were obtained from LI-COR, i.e., IRDye 680 Goat anti-Mouse (1:10,000, cat.#926-68070, Licor-Biosciences) or IRDye 800 Goat anti-Rabbit (1:10,000, cat.#926-32211, Licor-Biosciences). Proteins were detected on the Odyssey Infrared Imaging system (LI-COR Bioscience), and the analyses of variant proteins quantified by GFP were performed by the Image Studio Lite Ver 5.2 software. All full western blot images can be found in **Supplementary Fig 15**.

For enhanced chemiluminescence (ECL)-based detection, Immobilon Block-CH (Chemiluminescent Blocker) blocking buffer was used (cat.# WBAVDCH01, Millipore). The primary antibodies were diluted in this blocking buffer. The following secondary antibodies were used: Goat anti-mouse/HRP (1:10.000, cat.# A16078, ThermoFisher). Membranes were then incubated with Immobilon ECL Ultra Western HRP Substrate (Millipore) for 3-5 minutes and visualized using Amersham Imager 600 (GE Healthcare).

### CRISPR/Cas9 genome editing and clone screening

ZNRF3 truncating cells were generated via CRISPR-Cas9 genome editing. Different guide RNAs (gRNAs) (**Supplementary Table 6**), which were designed with the help of the CRISPR tool (http://crispor.tefor.net/), were inserted in the pSpCas9(BB)-2A-GFP (pX458) vector to introduce different *ZNRF3* truncating mutations in the genome of RNF43 deficient HEK293T cells (8). Approximately 40% confluent RNF43-KO HEK293T cells seeded into a 6-well plate were transfected with 600ng px458 using Lipofectamine 2000 following the manufacturer’s instruction. 48h after transfection, single GFP-expressing cells were sorted in 96-well plates by a fluorescence-activated cell sorter (FACS) FACSAria II cell sorter (BD Biosciences, San Jose, CA, USA). Two weeks later, a PCR screen was done to select the correct truncating mutation cells. The sequence alterations in ZNRF3 of RNF43-KO HEK293T clones are depicted in **Supplementary Fig 6**. All primers used for identifying sequence alterations can be found in **Supplementary Table 7**.

### Database analyses

The data reported in **Fig 1** and **Supplementary Fig 2** were obtained from the cBioPortal website by retrieving all ZNRF3 mutations from the curated set of non-redundant studies until May 2023. For the analyses reported in **Supplementary Table 8 and Fig 8**, we retrieved mutations in Wnt/β-catenin related genes specifically for all cancers carrying defective ZNRF3 variants or truncating mutations. As in most cases information about zygosity of the mutations was not available, this feature was not taken along in the analysis. For colorectal cancers we also retrieved BRAF mutation status, MSI-status and tumor location.

**Figure 1.**
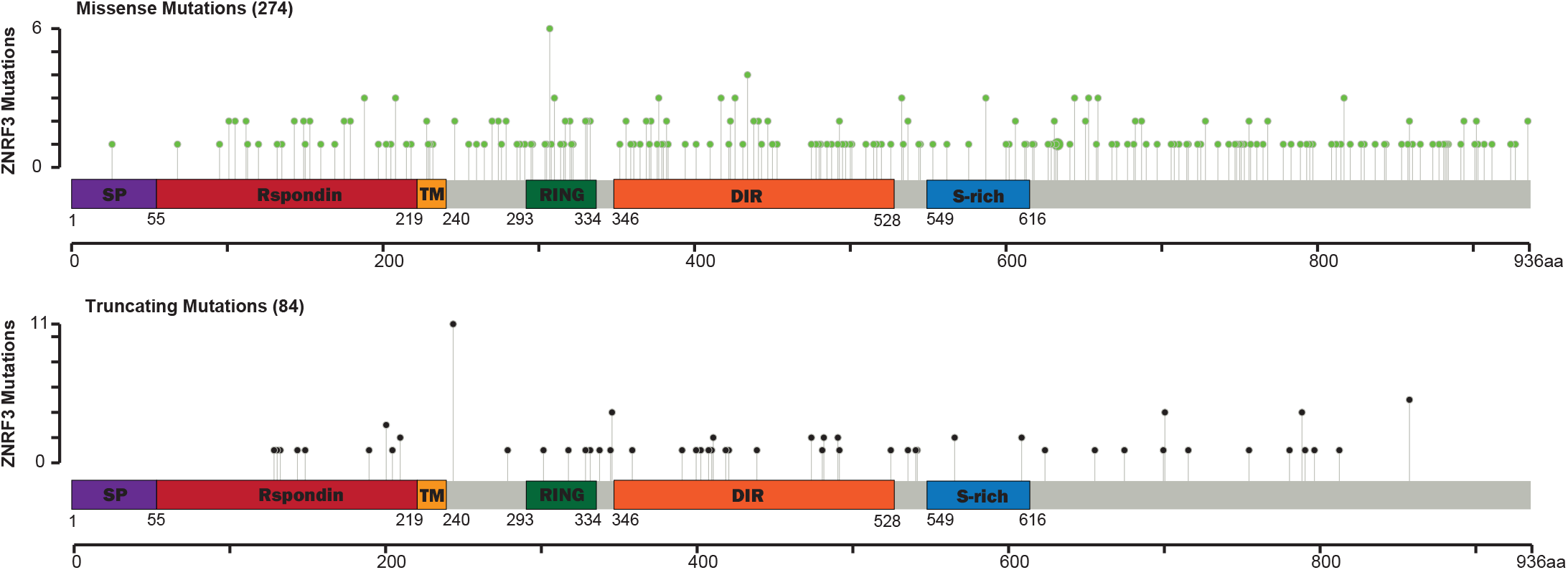
Schematic overview of the ZNRF3 protein and mutational landscape analysis. ZNRF3 missense and truncating mutations reported in the cBioPortal human cancer genome database. The number of the truncating mutations distributed along the entire ZNRF3 protein with a few hotspots is less than the missense mutations. The following protein domains are indicated: a signal peptide (aa 1-55), the extracellular R-spondin-binding domain (aa 56-219), a transmembrane domain (aa 220-240), the RING finger domain (aa 293-334), the disheveled interaction region (DIR) (aa 346-528) and a serine-rich region at the C-terminus (aa 549-616). The numbers correspond to the aa positions of the long ZNRF3 isoform.

### Statistical analysis

All results are presented as the mean ± standard deviation (SD). Statistical analyses were carried out using GraphPad Prism version 8.0.2 software (GraphPad Software Inc., San Diego, California, USA, RRID:SCR_002798). Differences were considered significant at a P value less than 0.05 (* P ≤ 0.05, ** P ≤ 0.01, *** P ≤ 0.001, **** P ≤ 0.0001).

## Results

### 1. Analysis of the ZNRF3 mutational landscape

We systematically examined the distribution of ZNRF3 mutations in multiple human cancer types using cBioPortal (http://www.cbioportal.org/). As shown in **Fig 1**, truncating and missense variants appear evenly distributed along the *ZNRF3* coding region, except for a lack of mutations in the first 100 amino acids encoded by exon 1. This latter exon is extremely GC-rich (**Supplementary Fig 1**), and for that reason will probably often be missed by next-generation sequencing approaches (14). Common germline ZNRF3 amino acid variants identified using the gnomAD database (https://gnomad.broadinstitute.org/) are not observed, meaning that basically all tumor-associated variants will be somatically acquired. Mutations are prevalent in uterus (5.8%) and bowel cancers (5%), with a predominance of missense mutations (**Supplementary Fig 2**). Other tumor types with ZNRF3 mutation frequencies above 1% are those of the skin, liver, lungs, and cancers of the esophagus/stomach.

### 2. The ZNRF3 long protein isoform is endogenously most relevant

Both a long ZNRF3 protein isoform (936aa) and shorter version (836aa) are reported in the Ensembl genome browser. The short isoform lacks the first 100 amino acids (**Supplementary Fig 3**). To investigate which isoform is most relevant endogenously, we generated vectors expressing both ZNRF3 long and short protein isoforms. A β-catenin reporter assay showed that the ZNRF3 long isoform can significantly inhibit Wnt-induced β-catenin signaling in HEK293T cells, while this is not the case for the short variant (**Supplementary Fig 3**). Next, we performed qPCR to test the expression of both *ZNRF3* RNA variants in colorectal (n=4), liver (n=8) and pancreatic cancer (n=4) cell lines. Although the short transcript can be detected in some cell lines, its expression is generally very low (**Supplementary Fig 4**). Last, using signal peptide prediction shows that only the long isoform contains a genuine signal peptide at its N-terminus (**Supplementary Fig 5**). Taken together, these observations show that only the ZNRF3 long isoform is clearly expressed and has the potential to transfer to the cell membrane to perform its principal function in reducing Wnt/β-catenin signaling.

### 3. Analysis of ZNRF3 truncating mutations

Various truncating *ZNRF3* mutations have been observed in cancers (**Fig 1**), but thus far their functional consequences are largely unknown. Therefore, we generated expression vectors that can be grouped into two types of truncations (**Fig 2A**). The first group delineates known functional domains (R246tr, S355tr, R536tr, R789tr), while the second set of truncations is potentially linked to a reported regulation of ZNRF3 by Caseine Kinase 1 (CK1). Both RNF43 and ZNRF3 are regulated in their function by CK1, which binds and phosphorylates residues in their S-rich domains (RNF43: ±455-507; ZNRF3: ±540-600) (15–17). For RNF43 it was shown that truncating mutations in a region (D504-Q563) directly following this S-rich domain, lead to an aberrant Wnt-independent increase in β-catenin signaling by trapping CK1 at the plasma membrane (17). Although ZNRF3 shows little homology with RNF43 in its C-terminal domain, we tested three tumor-associated truncating mutations reported in the equivalent ZNRF3 region (G612tr, G619tr, Q657tr).

**Figure 2.**
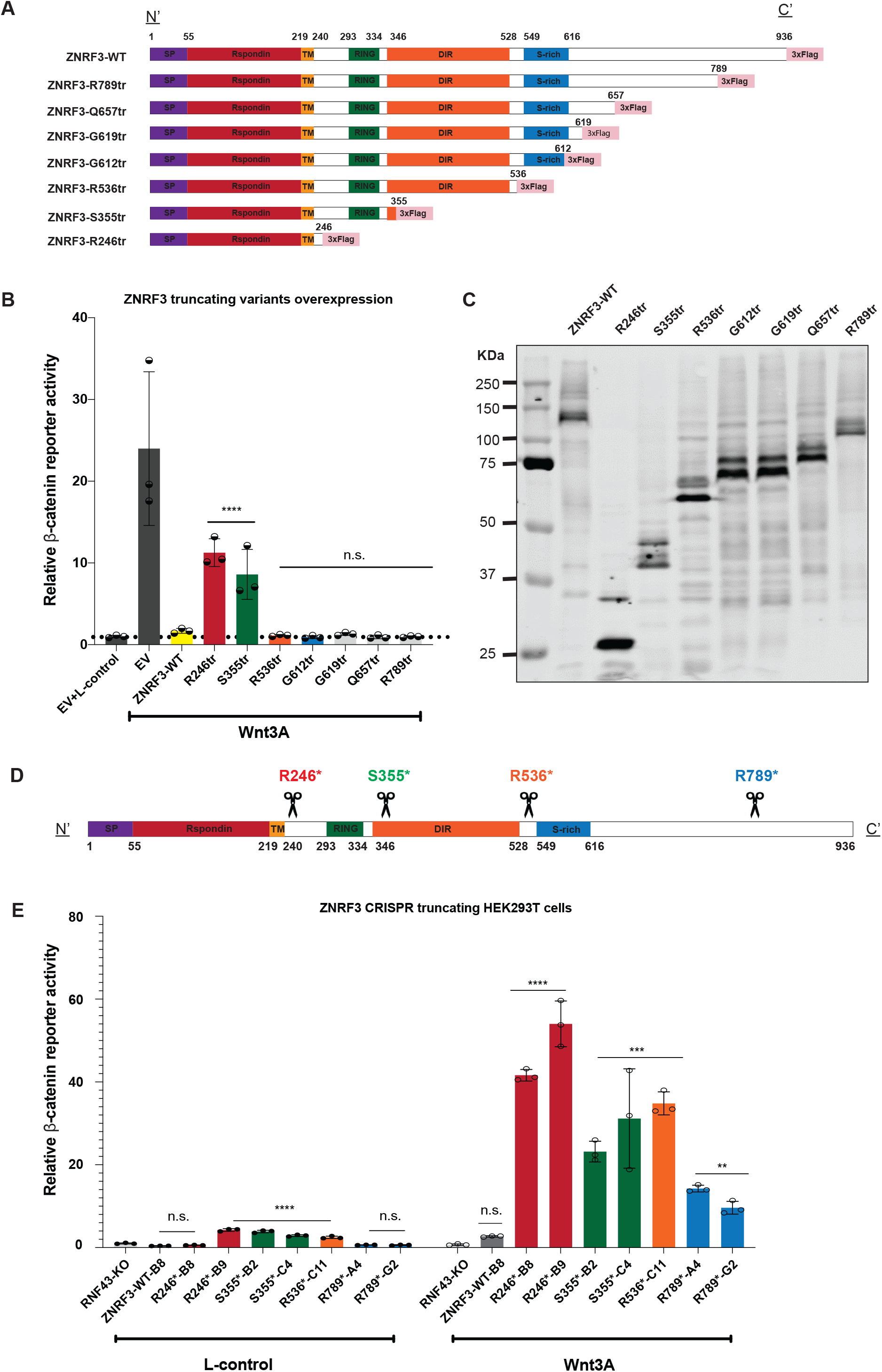
Characterization of ZNRF3 truncating mutations in regulating β-catenin signaling. All tested ZNRF3 truncations lose some regulatory functions at the endogenous level, while the longer variants retain partial activity. **(A)** A schematic diagram depicting the 3xFLAG-tagged ZNRF3 truncating mutant expression vectors that were generated. For clarity, we refer to each domain by reported ZNRF3 functional domains, except for the CK1 binding domain, which overlaps with the S-rich domain but has not yet been identified accurately. **(B)** A β-catenin reporter assay in HEK293T cells shows that the variants retaining at least the DIR were able to inhibit Wnt-induced β-catenin signaling comparable to wild-type ZNRF3. Substantial and significant increases in signaling are observed in the shorter variants R246tr and S355tr. The relative Wnt β-catenin signaling activities are depicted as WRE/CMV-Renilla ratios, in which the value obtained for wild-type (WT) ZNRF3 was arbitrarily set to 1. Wnt3A conditioned medium was added to Empty Vector (EV) and all ZNRF3 variants transfected cells. L-control medium was added to EV transfected cells as a negative control for β-catenin signaling. **(C)** Immunoblot showing correct expression of the expected variants following HEK293T transfection. **(D)** A schematic overview of the ZNRF3 truncation variants generated in HEK293T-RNF43-knockout (KO) cells using CRISPR/Cas9-mediated gene editing. The scissors locations represent the CRISPR/Cas9 cutting sites where frameshift mutations were introduced. **(E)** RNF43-KO cells were used to exclude the influence of homologous RNF43. ZNRF3-WT-B8 cells represent an additional ZNRF3-WT control clone lacking RNF43, obtained during the gene-editing process. The relative β-catenin reporter activities are depicted as WRE/CMV-Renilla ratios, in which the value obtained for ZNRF3-WT-B8 clone treated with L-control medium, was arbitrarily set to 1. When exposed to L-control conditioned medium, all truncations except R246*-B8, R789*A4, and R789*G2, show some loss of β-catenin regulatory function when expressed at endogenous levels. In the Wnt3A conditioned medium, all truncations show some loss of β-catenin regulatory function, while the longer variants retain more function. All Dual-luciferase data are shown as mean ± SD. Statistical significance was assessed using a one-way ANOVA test (\*\*\*\**P*<0.0001, *** *P*<0.001, ** *P*<0.01).

When expressed transiently in HEK293T cells, all variants retaining at least the Dishevelled interaction region (DIR) were able to inhibit Wnt-induced β-catenin signaling comparable to wild-type ZNRF3 (**Fig 2B/C**). Shorter variants lost most of their ability to inhibit β-catenin signaling. Unlike RNF43, truncations of ZNRF3 directly following the S-rich domain did not lead to a dominant activation of β-catenin signaling (17).

These results show that ZNRF3 and long variants thereof can effectively inhibit β-catenin signaling when overexpressed. To test this at endogenous levels we generated CRISPR-Cas9-mediated HEK293T clones carrying ZNRF3 truncations (**Fig 2D, Supplementary Fig 6**). RNF43 knockout cells were used to exclude the effect of RNF43 inhibiting β-catenin signaling (8). Two independent clones were obtained for each truncating variant in most cases. In contrast with the overexpression experiments, all truncations show some loss of β-catenin regulatory function when expressed at endogenous levels (**Fig 2E**). Loss of functionality is inversely correlated to the length of the truncated ZNRF3 protein, with the longest R789tr protein retaining most functionality. Truncations following the RING domain (S355tr) or DIR domain (R536tr) show intermediate defects. Taken together, our results indicate that at endogenous levels all tested truncations lose some regulatory function, while the longer variants retain partial activity.

### 4. Identification of missense ZNRF3 variants affecting **β**-catenin signaling

Currently, all reported ZNRF3 missense variations are regarded as variants with uncertain significance (VUS), meaning that it is unknown whether they affect ZNRF3 protein function. In total 274 unique tumor-associated variants have been reported in cBioPortal until July 2023. To better understand which of these may act as driver mutations in tumorigenesis, we systematically analyzed patient-derived ZNRF3 missense mutants. Previously, we reported that RNF43 missense variants that affect its function are restricted to the RING and extracellular R-spondin domains (9). As these are also among the most homologous domains between RNF43 and ZNRF3 **(Supplementary Fig 7)**, we included all ZNRF3 variants in these domains reported at the time of our analysis. In addition, we included several variants identified at evolutionary conserved amino acids **(Supplementary Fig 8)** reported in the transmembrane domain, the DIR and S-rich domain. The C-terminal domain of ZNRF3 (605–935) shows a low level of evolutionary conservation, making it unlikely that variants in this domain will affect ZNRF3 function.

Next, we tested all variants for their capability to inhibit Wnt-induced β-catenin signaling in HEK293T cells. In the R-spondin-binding domain 12 out of 47 missense variants showed a significant loss-of-function (LOF) (**Fig 3A**), even though for some specific variants the LOF was modest, e.g. G105C and A188V. Within the RING domain region 14 out of 27 tested missense variants showed a significant (partial) LOF or hyperactivating behavior (**Fig 4A**). Among the 8 variants located at other domains only the D545N variant showed a significant but very modest loss of β-catenin regulatory function (**Supplementary Fig 9**). In total, we identified 27 out of 82 tested variants that (partially) impaired ZNRF3 function.

**Figure 3.**
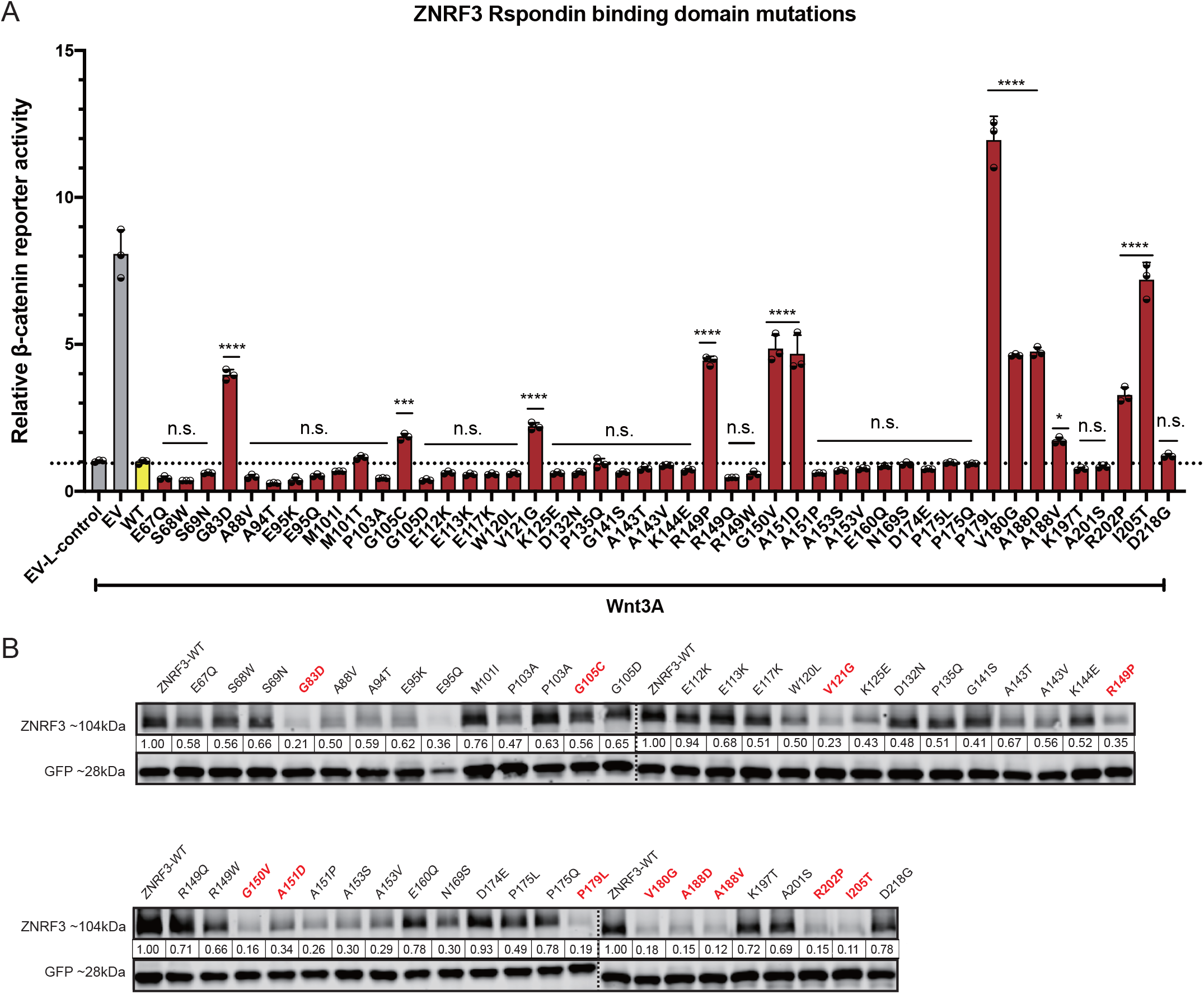
Analysis of missense variants within the ZNRF3 R-spondin-binding domain. Most variants in the ZNRF3 R-spondin-binding domain with (partial) loss of function show reduced protein stability. **(A)** A β-catenin reporter assay reveals 12 variants that show a significant loss of β-catenin regulatory activity. Wnt3A conditioned medium was added to Empty Vector (EV) and all ZNRF3 variant transfected cells. L-control conditioned medium was added to EV transfected cells as a negative control for β-catenin signaling. All relative β-catenin reporter activities are depicted as WRE/CMV-Renilla ratios, in which the value obtained for ZNRF3-WT was arbitrarily set to 1. Statistical significance was assessed using a one-way ANOVA test (\*\*\*\**P*<0.0001, *** *P*<0.001, * *P*<0.1). **(B)** All variants were co-transfected with an EGFP plasmid serving as transfection control. Using immunoblotting, 11/12 (partially) defective variants show a 3-9 fold reduction in protein stability, whereas most other variants retain normal or show a less reduced stability. All relative protein stabilities are depicted as ZNRF3/GFP ratios, in which the value obtained for ZNRF3-WT was set to 1.

**Figure 4.**
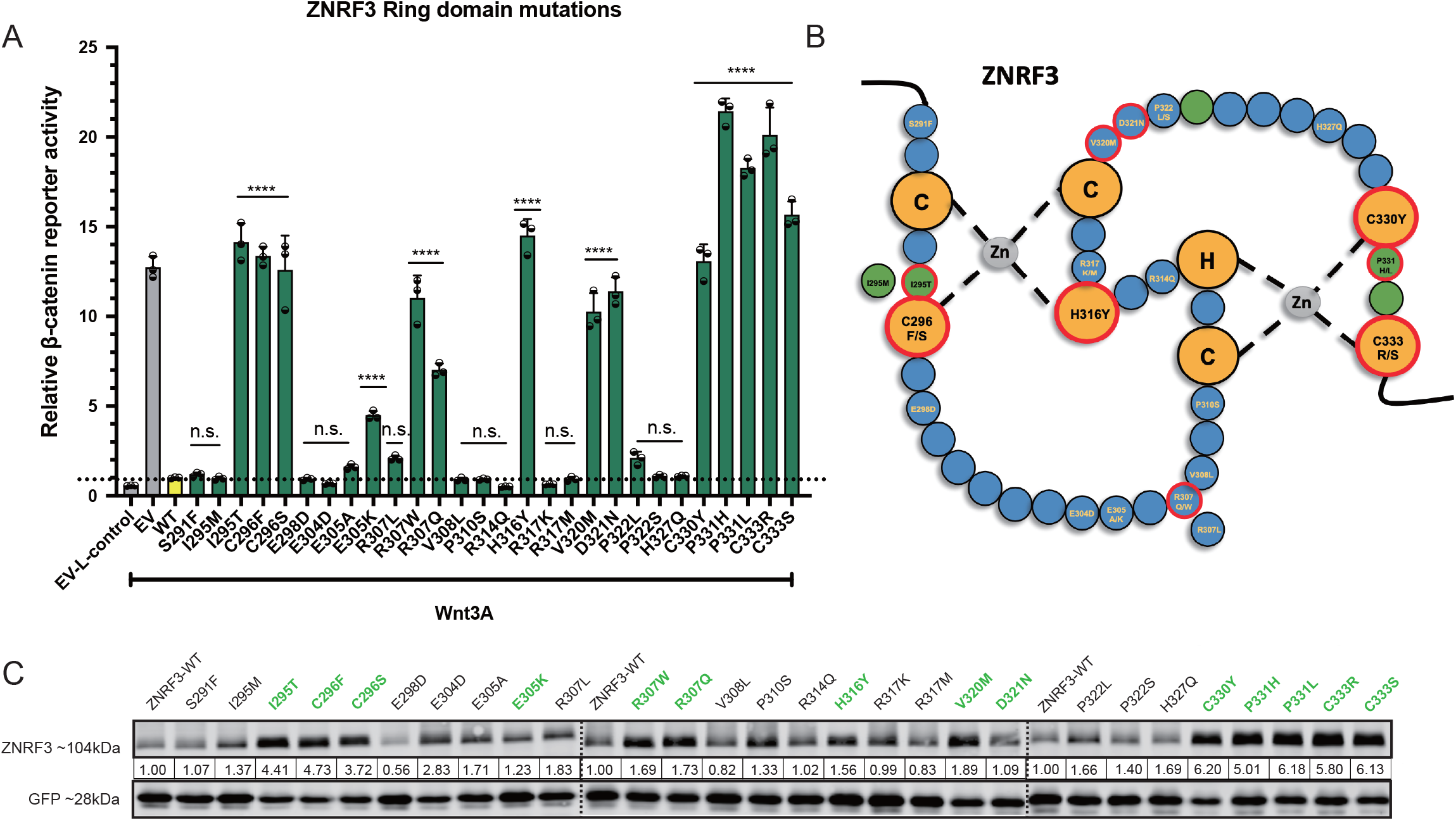
Analysis of missense variants within the ZNRF3 RING domain. About half of the tested RING domain variants result in a (partial) loss-of-function or hyperactivating behavior of ZNRF3. **(A)** A β-catenin reporter assay reveals 14/27 variants that show a significant loss of β-catenin regulatory activity. Wnt3A conditioned medium was added to Empty Vector (EV) and all ZNRF3 variants transfected cells. L-control conditioned medium was added to EV transfected cells as a negative control for β-catenin signaling. All relative β-catenin reporter activities are depicted as WRE/CMV-Renilla ratios, in which the value obtained for ZNRF3-WT was arbitrarily set to 1 (\*\*\*\**P*<0.0001). **(B)** Diagram depicting the RING domain structure of ZNRF3. The green circles are the residues required for interaction with ubiquitin-conjugating E2 enzymes, while yellow circles are the residues involved in Zinc (Zn) coordination. Tumor-associated variants that disrupt the β-catenin regulatory function of ZNRF3 are encircled by a red border. **(C)** All variants were co-transfected with EGFP plasmids. Immunoblot shows that all defective variants show at least a 1.5-fold increased protein stability, with the exception of E305K and D321N. All relative protein stabilities are depicted as ZNRF3 variants/GFP ratios, in which the value obtained from ZNRF3-WT was set to 1.

Focusing on the RING domain, as expected mutations of the C/H residues essential for Zinc interaction and formation of the RING structure, all disrupt β-catenin regulation (**Fig 4B**). The same holds true for mutants observed at residues required for interaction with ubiquitin conjugating E2 enzymes (green circles in **Fig 4B**) (18), with the exception of the I295M variant in which one hydrophobic residue is replaced by another one. Next, we made a comparison with reported variants of the homologous RING structure of RNF43 (9, 13), which we extended by investigating an additional 28 previously unexplored variants (**Supplementary Fig 10**). In total, 39 out of 44 RING domain variants disrupt RNF43 function, which is a significantly higher proportion than observed in the ZNRF3 RING domain structure (P<0.001, Chi-square test). **Supplementary Fig 11** provides schematic representations of the RING structures of RNF43/ZNRF3 and location of variants disrupting their function.

### 5. ZNRF3 variants defective in **β**-catenin regulation affect protein stability

To better define the mechanism through which LOF/hyperactivating ZNRF3 variants loose β-catenin regulatory activity, we first determined their protein stability. Basically all ZNRF3/RNF43 RING domain variants defective in β-catenin regulation, resulted in increased protein levels (**Fig 4C, Supplementary Fig 10B**), which is not unexpected given that these variants will also have lost their auto-ubiquitination potential (10, 11, 19).

The opposite effect was observed for 11 out of 12 R-spondin domain variants affecting β-catenin signaling (**Fig 3B**), which showed a clear 3-9 fold reduction in overall protein levels compared with wild-type ZNRF3. Similar observations were made for R-spondin domain variants within the RNF43 protein, but less consistently (**Supplementary Fig 12**). While several defective RNF43 variants show a 3-15 fold reduction in protein levels (A46V, I48T, P52L, G80R, P154L), others show a more modest about twofold reduction (G67C, A73F, A73V, A136T, P160S, A169T) or levels comparable to wild-type (K108E, G166C). Conversely, while most variants retaining normal β-catenin regulation in overexpression studies show protein levels equal to or at most twofold reduced compared to wild-type, the A78T variant is unsTable despite retaining β-catenin regulation. These results show that most defective R-spondin domain variants of ZNRF3 are reduced in protein stability, while this is the case for several but not all RNF43 variants.

### 6. LOF missense mutants in ZNRF3 R-spondin domain fail to reach the membrane correctly

We used a cell surface biotinylation assay to determine if the defective missense ZNRF3 variants can reach the plasma membrane correctly (**Fig 5A**). As shown in **Fig 5B**, all tested RING domain variants are clearly present at the membrane, and are detected at increased levels compared to wild-type ZNRF3, consistent with their increased stability. In contrast, defective variants within the R-spondin-binding domain are barely detecTable using this assay, indicating that they are strongly impaired in being transferred from the endoplasmic reticulum (ER) to the plasma membrane.

**Figure 5.**
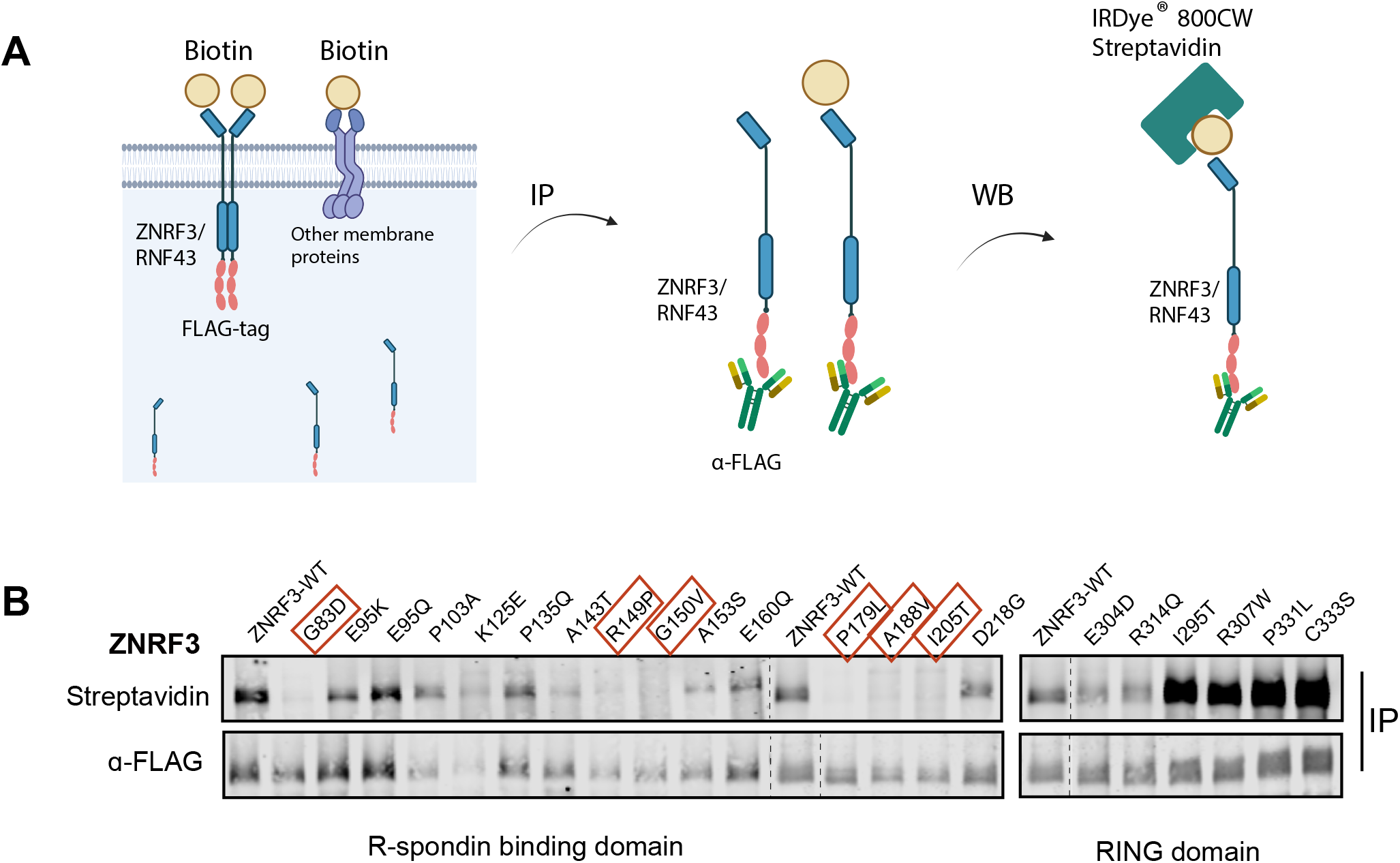
LOF missense mutants in ZNRF3 R-spondin domain fail to reach the membrane correctly. Analysis of plasma membrane presence of selected R-spondin and RING domain ZNRF3 variants. **(A)** Schematic overview showing the process of biotinylation and IP experiments. The Sulfo-NHS-LC-Biotin reagent is added to live cells for cell membrane protein labeling, followed by α-FLAG beads immunoprecipitation of ZNRF3/RNF43 FLAG-tagged protein. Next, IRDye800CW-streptavidin is used to label Biotin to measure the protein levels of ZNRF3/RNF43 reaching the cell membrane. **(B)** In contrast to wild-type ZNRF3 and non-defective R-spondin domain variants, six tested variants defective in β-catenin regulation (red squares) fail to reach the membrane correctly. Defective RING domain variants (I295T to C333S) have no problems in reaching the membrane and are detected at increased levels in line with their increased overall stability.

### 7. Low temperature partially rescues missense mutations in the R-spondin-binding domain

Both the reduced protein stability of R-spondin domain variants as well as their failure to reach the membrane, are indicative of a problem with protein misfolding. Within the endoplasmic reticulum, the structures of newly synthesized proteins undergo a rigorous quality control and when improperly folded are removed by ER-associated degradation (20). Similar observations have been made for other cell surface located proteins. One well-known example is the common F508del mutation in CFTR, leading to cystic fibrosis. The main consequence of this mutation is a defect in protein folding and trafficking, which can be rescued by growing cells at reduced temperatures, which has also been observed for other proteins (21, 22). Therefore, we tested if the membrane location of R-spondin variants can be restored by culturing cells at 27°C. We tested two variants for both RNF43 and ZNRF3 that all show reduced membrane presence at 37°C. In all four cases surface biotinylation was increased when cultured at lower temperature, although not reaching the levels of the wild-type proteins (**Fig 6A**). Next, using a β-catenin reporter assay we determined if low culture temperatures can also significantly affect β-catenin regulation. Out of 12 tested ZNRF3 variants, 7 showed a near-complete restoration of their regulatory function, while this was only observed for 2 out of 14 RNF43 variants (**Fig 6B/C)**. Taken together, our results indicate that the main functional consequence of R-spondin domain mutants of especially ZNRF3, is a failure to fold correctly, leading to reduced protein stability and trafficking to the membrane. These defects are temperature-sensitive as they can partially be rescued by lower temperatures.

**Figure 6.**
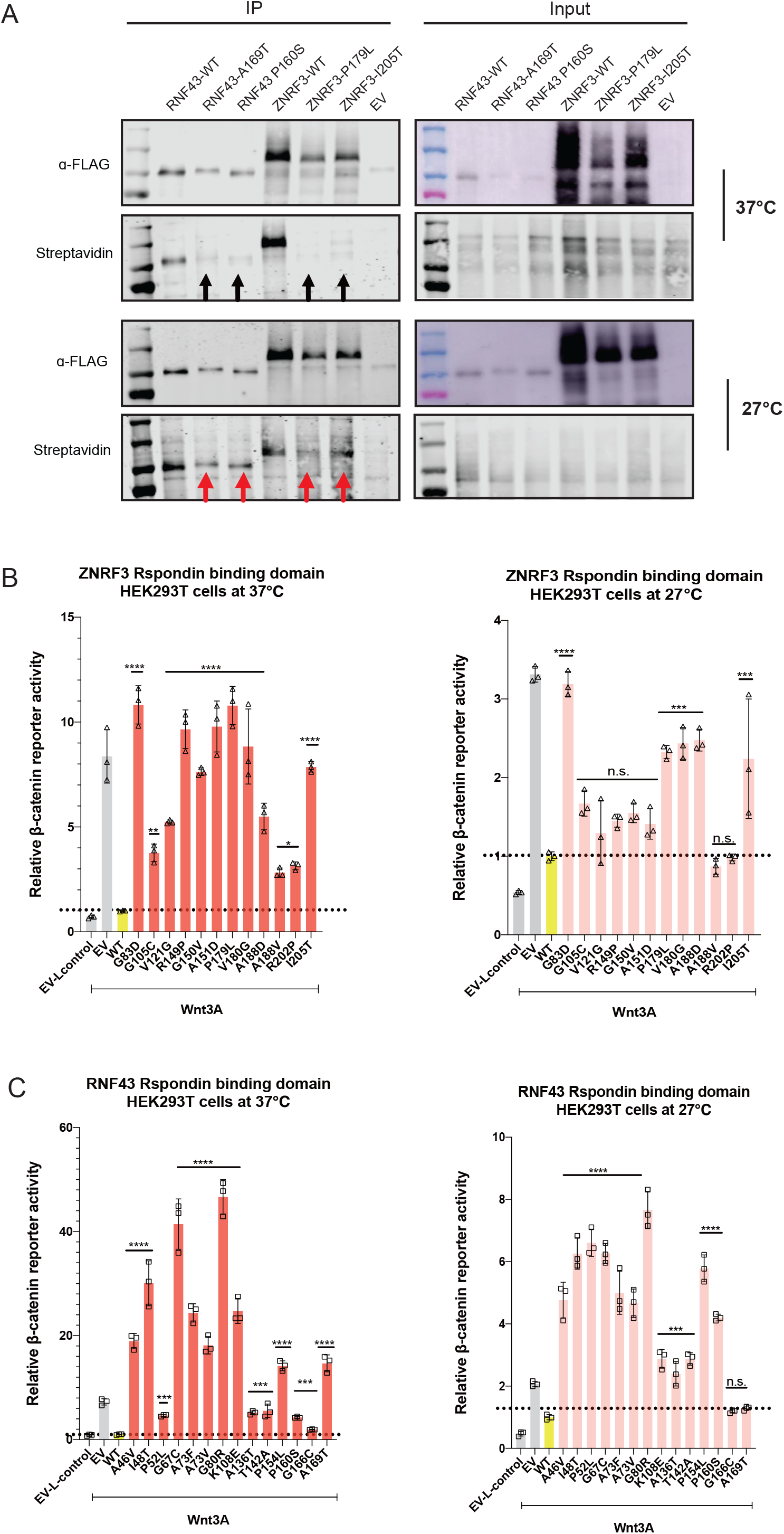
Low temperature partially rescues missense mutations in the R-spondin-binding domain. Culturing cells at 27°C supports variants of the ZNRF3 R-spondin-binding domain to (partially) regain their function in regulating β-catenin signaling. **(A)** Left panel shows an IP-experiment of surface-biotinylated RNF43 and ZNRF3 R-spondin domain mutants cultured at 37°C or 27°C. Mutants fail to reach the cell membrane when cultured at 37°C, while they show improved membrane presence when grown at 27°C. **(B)** An assessment of β-catenin reporter activity shows that 7/12 variants in the ZNRF3 R-spondin-binding domain regain functionality in Wnt/β-catenin signaling regulation when cultured at 27°C. **(C)** The same analysis performed for 14 RNF43 variants shows that only the G166C and A169T variants can be rescued at 27°C. Wnt3A conditioned medium was added to Empty Vector (EV) and all RNF43/ZNRF3 variants transfected cells. L-control conditioned medium was added to EV-transfected cells as a negative control for β-catenin signaling. All relative β-catenin reporter activities are depicted as WRE/CMV-Renilla ratios, in which the value obtained for RNF43-WT or ZNRF3-WT was arbitrarily set to 1. Statistical significance was assessed using a one-way ANOVA test (****P<0.0001, *** P<0.001, * P<0.1).

### 8. Testing “dominant-negative” potential of R-spondin/RING variants at endogenous levels

Already in the first papers linking RNF43 and ZNRF3 to Wnt/β-catenin signaling, it was noted that mutating their RING domains led to hyperactivation of β-catenin signaling, indicating a dominant-negative mode of action (10, 11). In subsequent years this observation was confirmed in several publications and also extended to mutations in the R-Spondin domain (9, 12, 13). Here, we show the same for specific ZNRF3 variants (e.g. P179L, P331H). The underlying mechanism relies on the ability of RNF43 and ZNRF3 to form homo- and heterodimers (9, 23). It is believed that if one of the dimer-partners carries an inactivating RING or R-spondin mutation, it can render the entire dimer inactive in a dominant-negative manner (**Supplementary Fig 14A)**. This would explain why transient overexpression of such variants leads to hyperactivation, as it will also inhibit endogenously expressed RNF43/ZNRF3 (9). This observation has led to the speculation that these missense variants may have oncogenic potential (9), meaning that one such RNF43/ZNRF3 mutation may already sufficiently increase β-catenin signaling to drive tumorigenesis, and that no second hit is required.

Thus far this speculation is entirely based on overexpression experiments, which, however, leads to a more than 1000-fold overexpression on RNA level (**Supplementary Fig 13A**), and most likely also on protein level. Therefore, we decided to test the relevance of “dominant-negative” variants at endogenous levels. Using gene-editing we generated various HEK293T clones expressing R-spondin (P179L) or RING domain ZNRF3 variants (R307W, P311L/H) in a heterozygous fashion. On purpose, we used wild-type HEK293T cells to determine if the remaining wild-type RNF43 and ZNRF3 can be sufficiently inhibited. HEK293T cells express *ZNRF3* at approximately 20-30-fold higher levels than *RNF43* (**Supplementary Fig 13B**) (10), indicating that expression levels of mutant ZNRF3 are likely sufficiently high to impose a dominant-negative effect on wild-type RNF43/ZNRF3. Importantly, we also generated four clones carrying a heterozygous *ZNRF3* knockout, to allow a direct comparison between loss-of-function and supposed dominant-negative mutations.

Next, we exposed all clones to exogenous Wnt3A and determined β-catenin reporter activity (**Fig 7A, Supplementary Fig 14B**). Although we do observe some clone-to-clone variation, basically all heterozygous mutant clones show a modest 2-8 fold activation of β-catenin signaling compared to wild-type cells. Importantly, we see no clear difference in activation when comparing heterozygous knock-out of ZNRF3 (lane 5) with clones heterozygously expressing “hyperactive” variants of ZNRF3 (lanes 2-4), indicating that the proposed dominant-negative behavior does not exist or is very weak. The low level induction of β-catenin signaling in all heterozygous clones, also suggests that partial loss of ZNRF3 expression leads to haploinsufficiency, at least in HEK293T cells.

**Figure 7.**
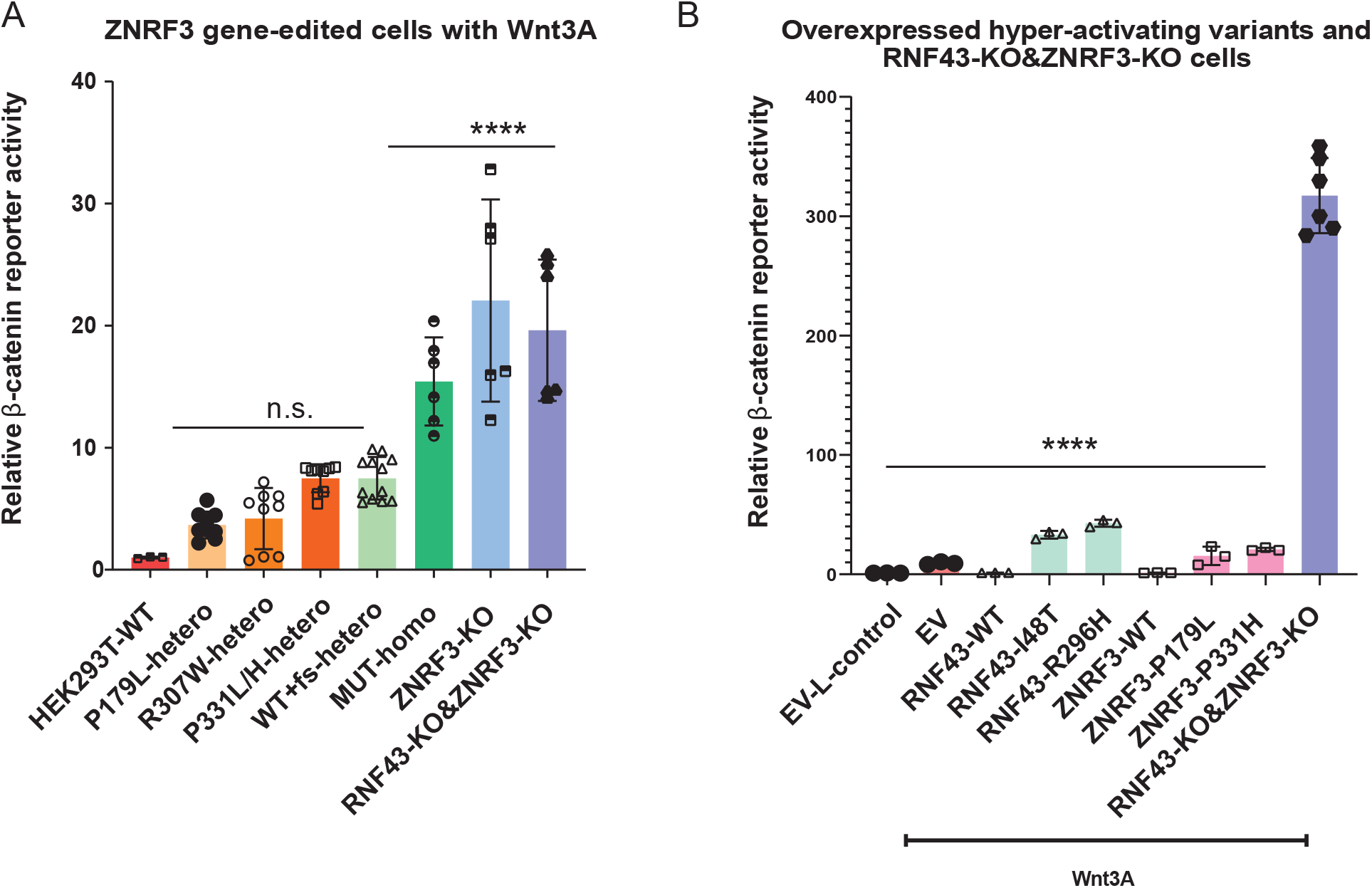
Testing dominant-negative potential of ZNRF3 variants at endogenous levels. Endogenous levels of heterozygous missense mutations do not affect ZNRF3 function more strongly than heterozygous knockout mutations. β-catenin reporter assays for knockin/out ZNRF3 clones. All heterozygous mutations, including P179L, R307W, P331L/H, and WT+frameshift (fs), slightly increase β-catenin signaling (2-8 fold) to similar levels. Homozygous ZNRF3 knockout clones, homozygous R307W or P331H knockin clones, and RNF43/ZNRF3 double knockout clones, all show a comparable further increase in signaling (18-30 fold). The relative Wnt/β-catenin signaling activities are depicted as WRE/CMV-Renilla ratios, in which the value obtained for wild-type (WT) HEK293T cells was arbitrarily set to 1. Wnt3A conditioned medium was added to WT HEK293T cells and all CRISPR-edited HEK293T cells. Statistical significance was assessed using a T-test (****P<0.0001, ** P<0.01, * P<0.1). For each heterozygous amino acid variant, 3-4 individual clones were used. Values for each individual clone are shown in Supplemental Figure 14B. **(B)** A β-catenin reporter activity analysis performed for representative supposedly dominant-negative RNF43 or ZNRF3 variants, shows that overexpression of these hyperactive variants reduces β-catenin signaling much weaker than a complete knockout of both proteins. Wnt3A was added to all samples except for control-treated empty vector (EV) transfected cells, serving as negative control. All relative β-catenin reporter activities are depicted as WRE/CMV-Renilla ratios, in which the value obtained for EV with L-Control was arbitrarily set to 1. Statistical significance was assessed using a one-way ANOVA test (****P<0.0001).

**Figure 8.**
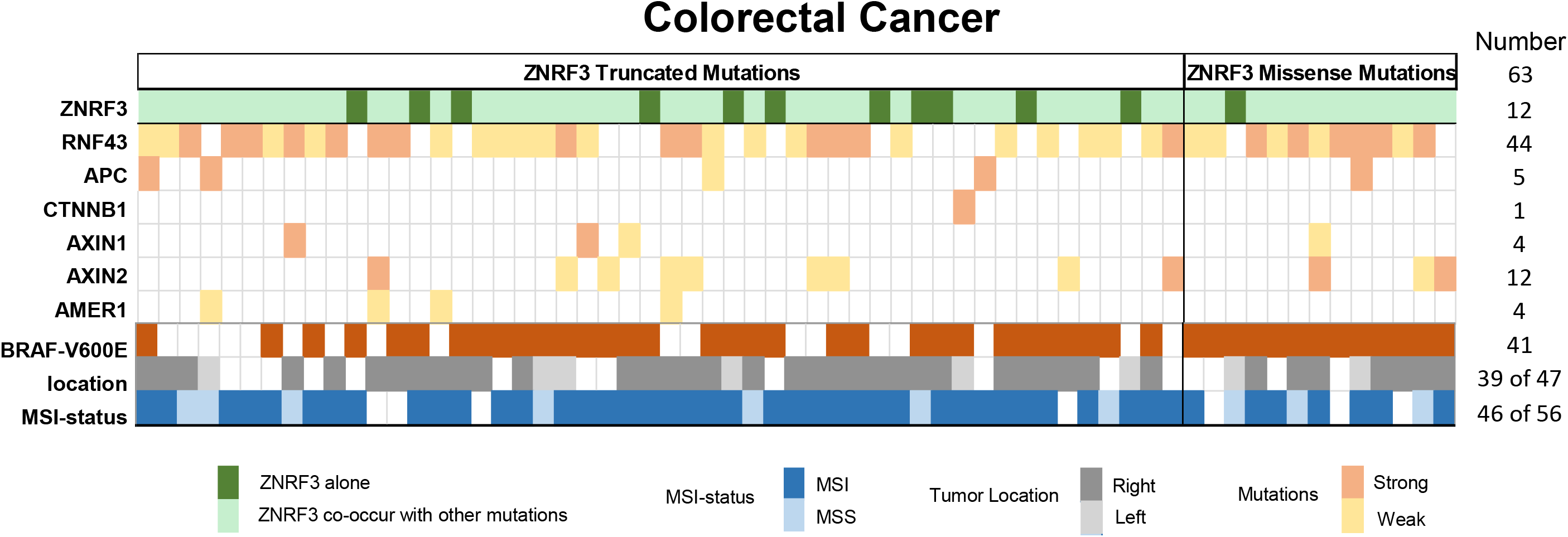
Co-occurrence with other β-catenin activating mutations in ZNRF3-mutant colorectal cancers. From the cBioPortal database, we obtained information from ZNRF3-mutant colorectal tumors about APC, CTNNB1, AXIN1, AXIN2, RNF43 and BRAF mutation status, in addition to CRC tumor location and MSI-status. Only tumors with missense ZNRF3 variants shown to be defective in our analysis were included, in addition to all CRCs carrying truncating ZNRF3 mutations. CRCs only carrying ZNRF3 mutations are indicated with a dark-green color. For each additional β-catenin related gene, we have indicated whether the mutation is expected to strongly impair protein function or only partially, using a dark-orange or light-golden color coding, respectively. A more extensive description for the criteria used to generate this Figure is provided in **Supplemental Table 8**.

We also obtained clones with homozygous expression of R307W and P331H variants, both of which exhibit increased β-catenin reporter activity compared to heterozygous clones (**Fig 7A, Supplementary Fig 14B**). However, in a direct comparison with previously generated ZNRF3- and double RNF43/ZNRF3 knockout clones, no significant difference is observed.

To further demonstrate that the proposed dominant-negative behavior of hyperactive variants is rather weak, we performed a direct comparison of β-catenin reporter activity observed in double RNF43/ZNRF3 knockout clones and those of overexpressed hyperactive variants. If overexpression of these variants would be very effective in blocking endogenous RNF43/ZNRF3, it would result in a comparable activation of β-catenin signaling. However, induction of β-catenin signaling by selected RNF43 and ZNRF3 hyperactive variants is much lower compared with a complete knockout of both proteins (**Fig 7B**).

To summarize, at endogenous expression levels all tested RING and R-spondin ZNRF3 domain variants, do not possess sufficient dominant-negative activity to impair the function of the remaining wild-type RNF43 and ZNRF3 proteins. In other words, they appear to behave as classical loss-of-function mutations.

### 9. ZNRF3 mutations often co-occur with other **β**-catenin activating gene mutations in colorectal cancer

Thus far our data show that ZNRF3 truncating mutations can be regarded as (partial) loss of function mutations, while this applies to a selected number of missense mutations. We explored the cBioPortal database to identify which cancers carry these defective ZNRF3 variants (**Supplementary Table 8**). They are identified at low numbers in various cancer types, such as those of the endometrium, stomach, liver and adrenal gland, but are most frequently observed in colorectal cancers (63/7435 CRC patients, i.e. 0.85%). As shown in **Fig 8** and **Supplementary Table 8**, CRCs with defective ZNRF3 mutations are highly enriched for BRAF-V600E mutation, a mismatch-repair defect, and right-sided tumor location (41/63, 46/56, and 39/47, respectively). Next, we determined to what extent *ZNRF3* mutations are accompanied by other gene mutations that result in β-catenin activation (**Fig 8, Supplementary Table 8**). Out of 63 cancers, we identified 12 that apparently only carry inactivating *ZNRF3* mutations. ZNRF3-mutant CRCs carrying additional *APC* truncations or oncogenic *CTNNB1* mutations classically linked to β-catenin activation, are only observed in 5 and 1 cancer, respectively. The remaining 45 cancers carry additional truncating mutations in either *AXIN1*, *AXIN2* or *AMER1*, or defective missense or truncating *RNF43* mutations (8, 9, 24, 25). Especially, the characteristic mismatch repair defect associated G659Vfs*41 truncating RNF43 mutation resulting in a partially defective protein (8, 24), is observed in a large fraction of tumors (31/63). Thus, in colorectal cancer *ZNRF3* mutations are often accompanied by other gene mutations that are expected to weakly increase β-catenin signaling.

## Discussion

The *ZNRF3* and *RNF43* genes encode for homologous transmembrane E3-ubiquitin ligases that remove Wnt-receptors from the membrane, thereby limiting the level of Wnt-induced β-catenin signaling (2–5). They are frequently mutated in various cancer types resulting in increased β-catenin signaling, which, importantly, is dependent on extracellular Wnt ligand exposure (2). This latter feature has led to an extensive research to identify therapeutic options that reduce extracellular Wnt ligand binding to tumor cells, as RNF43/ZNRF3-defective cancers are likely to benefit from such treatments (6, 7). For these therapies to be effective, it must be known which RNF43/ZNRF3 mutations lead to sufficient loss of functional activity. While this has been explored in detail for RNF43 (8, 9, 12, 13, 17), it has been largely neglected for ZNRF3. To fill this knowledge gap, we extensively studied the functional consequences of truncating and 82 tumor-associated missense ZNRF3 variants. In addition, we have better defined the mode of action through which ZNRF3 and RNF43 mutations contribute to cancer formation, including at endogenous expression levels.

For truncating ZNRF3 mutations, all investigated variants partially lost β-catenin regulatory function at endogenous expression levels. An inverse correlation was observed with longer proteins retaining more function, but even the longest variant tested, i.e. R789tr, showed some loss of activity. This is reminiscent of RNF43, in which the most common G659fs mutation retains a substantial level of β-catenin regulation, but not as effective as wild-type RNF43 (8, 9). Currently, no clear function has been attributed to the C-terminal regions of either RNF43 or ZNRF3. In this domain, both proteins show little homology to each other and other known protein structures, and generally have a low level of evolutionary conservation. This lack of information makes it difficult to ascertain why loss of the C-terminus affects their function.

Specifically for RNF43, truncating mutations have been identified directly following the S-rich domain that trap CK1 kinase at the membrane, resulting in a Wnt-independent β-catenin activation (17). Although the exact mechanism is unknown and it involves only a small number of cancers (n=15 to date), it nevertheless represents an attractive alternative mechanism to aberrantly increase β-catenin signaling. However, ZNRF3 truncations at equivalent positions failed to affect β-catenin signaling, suggesting that the CK1 trapping phenomenon is restricted to RNF43.

The RING domain structure of RNF43/ZNRF3 represents its most critical functional domain, and shows more than 70% amino acid identity between both proteins. For RNF43, basically every tumor-associated variant in the RING domain disrupts its function. In contrast, ZNRF3 appears to be more tolerant for amino acid changes in the RING domain, as only half of the tested variants lead to a functional impairment. The main function of the RING domain is to associate with one of the approximately 30 E2 ubiquitin conjugating enzymes (18), which in turn then transfer ubiquitin to RNF43/ZNRF3 substrates, e.g. the Wnt-receptor. Which E2 enzymes effectively bind to RNF43 has only been sparsely investigated in yeast-two-hybrid assays or in vitro association studies (19, 26), and even less so for ZNRF3 (26), but our results suggest that the ZNRF3 RING structure is less rigid in finding an E2 binding partner.

The second main domain in which amino acid variants lead to loss of ZNRF3/RNF43 functionality, is the extracellular R-spondin-binding domain. The exact mechanism leading to their functional impairment is still not fully elucidated, but studies conducted prior to our analysis hinted at problems with correct trafficking of mutant proteins to the plasma membrane. For example, Jiang et al. showed that cell surface levels of an RNF43 F69C mutant are reduced compared to wild-type (27). Others reported mislocalization of RNF43 R-spondin-binding domain variants within the endoplasmic reticulum (9, 12). It has also been speculated that defective R-spondin domain variants may have reduced affinity for FZD or other membrane proteins (9). Here, we show that all six ZNRF3 mutants tested fail to reach the membrane correctly, which is accompanied by a 3-9 fold reduction in protein stability for 11/12 defective variants. Interestingly, membrane presence and functionality of several variants can be improved by culturing cells at reduced temperatures. All these findings are consistent with a defect in ZNRF3 protein folding and increased degradation through ER-associated degradation (20–22). Although RNF43 was not the main focus of our study, several but not all defective variants show reduced stability or fail to reach the membrane correctly (A169T, P160S). However, functional rescue at 27°C is rarely observed for RNF43 variants and the effects on stability are also less consistent compared to ZNRF3. Thus, it remains to be seen whether a defect in folding and trafficking represents a general explanation for the defective nature of RNF43 R-spondin domain variants.

A second main finding of our work is that the proposed hyperactivating/dominant-negative nature of many defective missense variants observed in transient overexpression experiments (9–12), is not relevant at endogenous levels. When transiently overexpressed, specific RNF43/ZNRF3 variants increase β-catenin reporter activity an additional 1.5-3 fold compared to adding Wnt3A alone. Mechanistically this was explained by assuming that mutant RNF43/ZNRF3 forms a dimer with wild-type proteins, rendering the entire dimer inactive. If correct, it would suggest that these variants have oncogenic potential, meaning that one such RNF43/ZNRF3 mutation may already sufficiently increase β-catenin signaling to drive tumorigenesis, and that no second hit is required. However, our results suggest that this is not the case for the following reasons. First, overexpression results in a more than 1000-fold increase at RNA levels, which likely also translates into a similar overexpression on protein level, even though we cannot faithfully compare with endogenous levels because of lack of specific antibodies (28). This implies that in the overexpression experiments basically all dimers formed will have incorporated a mutant RNF43/ZNRF3. If the dominant-negative behavior would be very effective, it would lead to an equivalent β-catenin activation as a full knockout of both RNF43 and ZNRF3, which is clearly not the case (**Fig 7B**). More importantly, we also confirm this at endogenous expression levels for *ZNRF3*. All clones generated to heterozygously express mutant ZNRF3 variants, show a comparable or even lower induction of β-catenin signaling when compared to heterozygous knock-out clones, while higher signaling levels would be expected in case of dominant-negative activity. Moreover, dimer formation becomes less likely for R-spondin domain variants leading to unsTable RNF43/ZNRF3 proteins, as there will be fewer mutant proteins to form a dimer. Based on these observations, we consider the supposed dominant-negative activity to be of minor relevance endogenously. We show that here clearly for ZNRF3, but based on the relatively weak induction when overexpressed, we also consider it likely for so-called hyperactivating RNF43 variants. In other words, at endogenous levels all RNF43/ZNRF3 variants appear to behave as classical loss-of-function mutations.

Mutations leading to a defective ZNRF3 protein were identified in several tumor types, but most frequently in colorectal cancers enriched for a mismatch repair defect, BRAF-V600E mutation and right-sided tumor location. The vast majority of CRCs acquire mutations that aberrantly increase β-catenin signaling (1, 29). Right-sided cancers generally select for mutations that lead to a lower level of β-catenin activation than their left-sided counterparts (29–32). This can be accomplished by obtaining longer APC truncations that retain more β-catenin binding- and breakdown repeats (29), but can also result from gene mutations that are expected to lead to more modest activation of β-catenin signaling. For example, it has been shown that inactivating AXIN1 mutations lead to only weak activation of β-catenin signaling due to partial compensation by AXIN2 and vice versa (25). The same applies to RNF43 and ZNRF3, which functionally compensate each other. This has been demonstrated, among others, by knockout studies in mice. A single knockout of either *Rnf43* or *Znrf3* does not lead to a discernable intestinal phenotype, while their simultaneous loss was required to observe a clear expansion of the proliferative compartment (11). Similarly, intestinal organoids can only grow independent of R-spondin when both genes are inactivated (33). These compensatory mechanisms can explain why most *ZNRF3* mutant CRCs acquire mutations at other genes that lead to a modest activation of β-catenin signaling, such as truncating AXIN1/AXIN2/AMER1 mutations or missense/truncating RNF43 mutations. Combined these weak activating mutations may increase β-catenin signaling to sufficiently high levels to support tumor growth on this side of the colon. A similar explanation may also apply to ZNRF3-mutant cancers of the endometrium and liver that also often show co-occurrence with other β-catenin activating mutations (**Supplementary Table 8**), while in other cancers such as melanomas and adrenocortical carcinomas the ZNRF3 mutation may be sufficient in itself.

To summarize, we extensively analyze tumor-associated ZNRF3 mutations and show that at endogenous levels, all truncating mutations tested exhibit loss-of-function, however, an inverse correlation is observed with longer variants retaining residual functionality. Several missense mutations were identified within the RING and extracellular R-Spondin-binding domain structures that lead to a (partial) loss-of-function/hyperactivation when overexpressed. Mechanistically, we show that for the ZNRF3 R-spondin domain mutants this results from reduced protein stability and a defect in trafficking to the membrane, which, interestingly, for some variants can be partially rescued by culturing cells at lower temperatures. These observations are in line with a protein folding defect and ER-associated degradation of mutant proteins. Similar observations were made for some, but not all RNF43 R-spondin domain variants. Importantly, we show that the dominant-negative hyperactivating nature of ZNRF3/RNF43 defective missense variants proposed previously, is not relevant at endogenous expression levels, meaning that these variants should be regarded as classical loss-of-function mutations. Finally, we show that defective ZNRF3 variants are most common in mismatch repair-defective colorectal cancers, where they co-occur with other gene mutations expected to lead to modest β-catenin activation.

## Acknowledgements

This research was financially supported by a China Scholarship Council PhD fellowship to Shanshan Li (File NO. 201909370083), Jiahui Niu (File NO. 202007660001) and Ruyi Zhang (File NO. 201808530490).

## Author contributions

SL, JN, SM, RZ, JvN, NdS, LvdK performed the experimental work as well as data analysis. SL authored the manuscript. MPP supervised the project. RS coordinated the project and authored the manuscript. All authors reviewed the results and approved the final version of the manuscript.

