## Supplemental Figures for "Lack of dominant-negative activity for tumor-associated ZNRF3 missense mutations at endogenous expression levels"

### Supplementary Fig.1

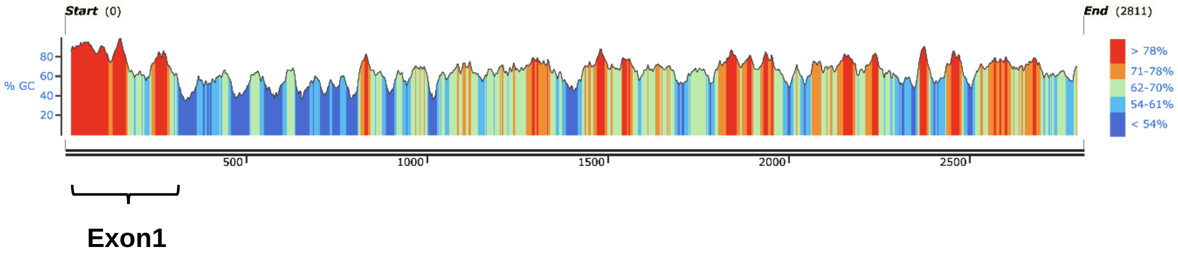


**Supplementary Figure 1.** Analysis of the GC content of the *ZNRF3* open-reading frame encoding the long isoform. The map was generated using Snapgene and shows that the GC content of the exon1 region is exceptionally high (>78%), which is challenging to amplify by PCR and detect mutations by sequencing.

### Supplementary Fig.2 page1

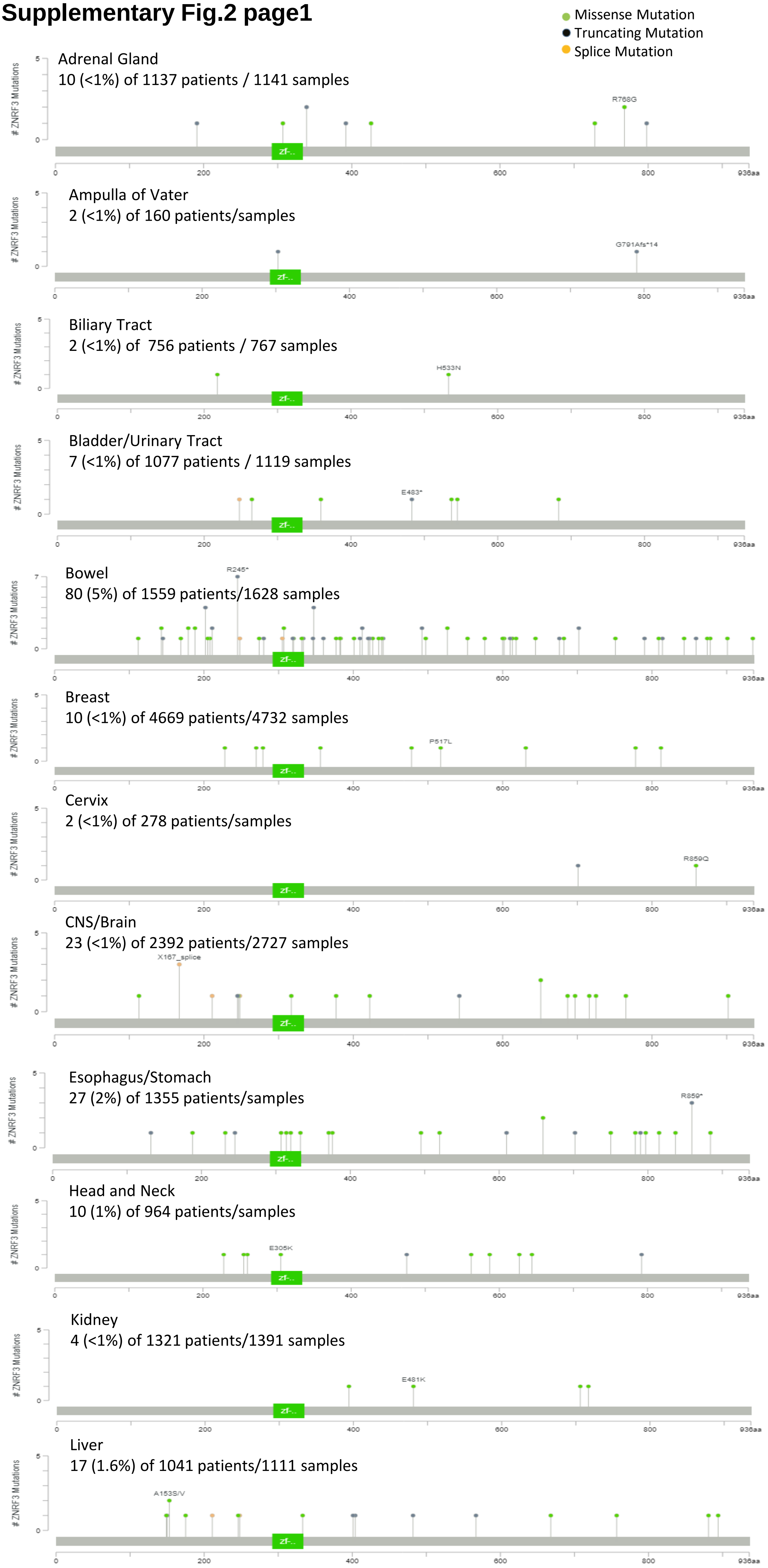

#### Supplementary Fig.2 page2

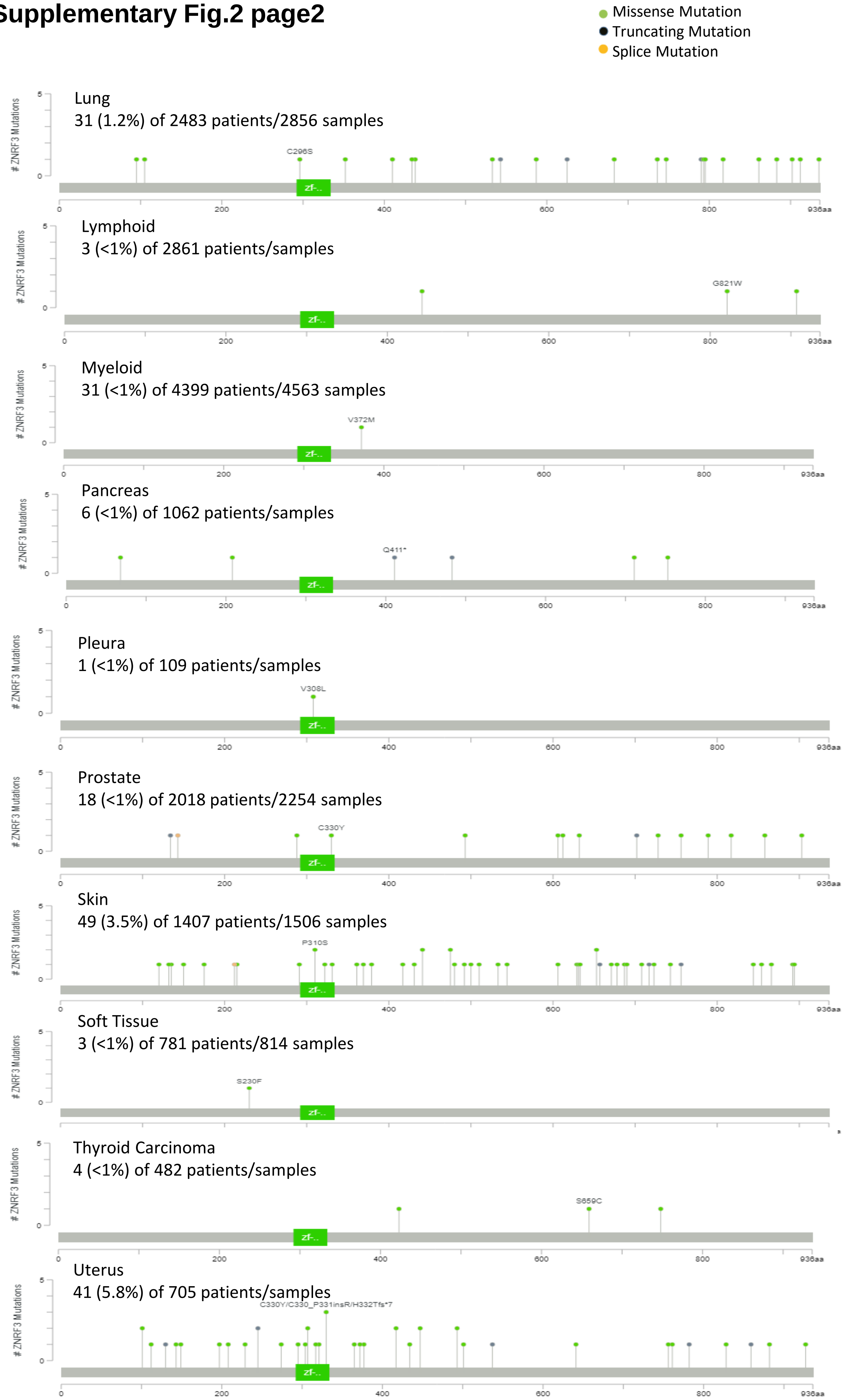


**Supplementary Figure 2.** The ZNRF3 mutation landscape for various cancer types was obtained from the cBioPortal database (http://www.cbioportal.org/). Data are updated until May 2023.

### Supplementary Fig.3

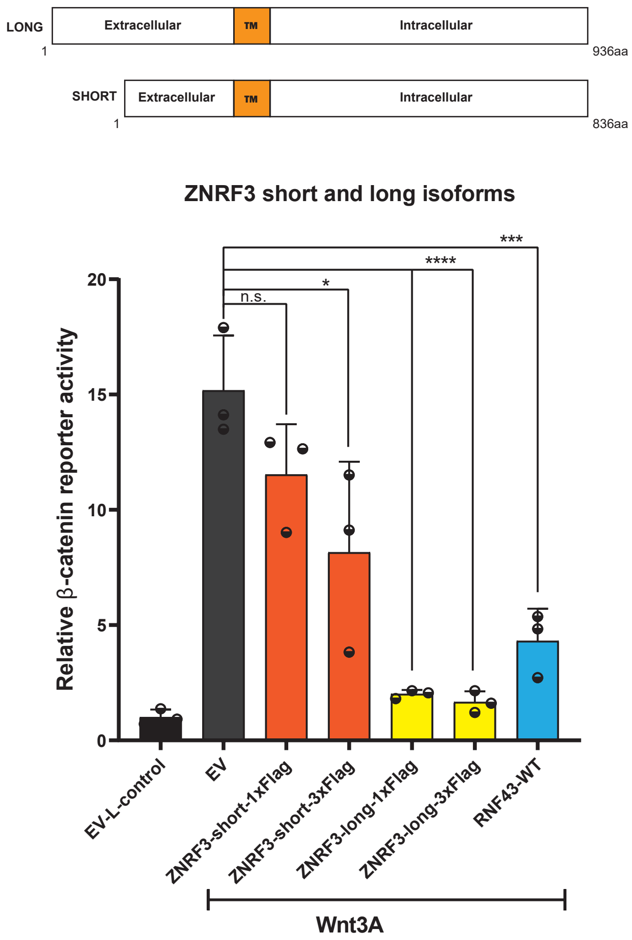


**Supplementary Figure 3.** A β-catenin reporter assay shows that the long ZNRF3 isoform is a stronger inhibitor of Wnt/β-catenin signaling. The plasmids that were generated with the coding sequence for the long and short ZNRF3 isoforms with 1x or 3xFlag-tagged expression were transfected into HEK293T cells. The relative β-catenin reporter activities are depicted as WRE/CMV-Renilla ratios, in which the value obtained for EV-transfected was arbitrarily set to 1. Wnt3A conditioned medium was added to Empty Vector (EV) and all ZNRF3 variants transfected cells. L-control medium was added to EV-transfected cells as a negative control for β-catenin signaling. Statistical significance was assessed using a one-way ANOVA test (****P<0.0001, *** P<0.001, *P<0.1).

**Supplementary Fig.4 page1**

### Colon Adenocarcinoma

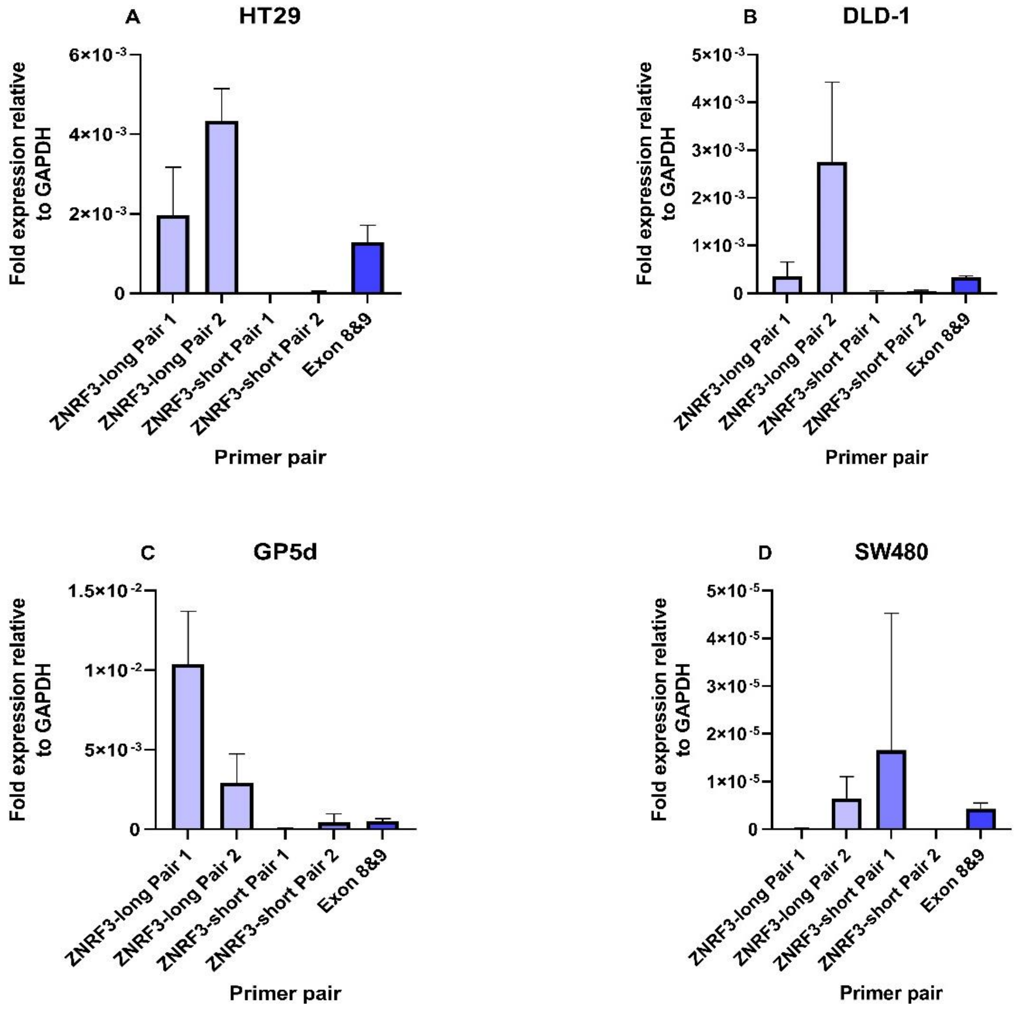


### Hepatoblastoma/Hepatocellular Carcinoma

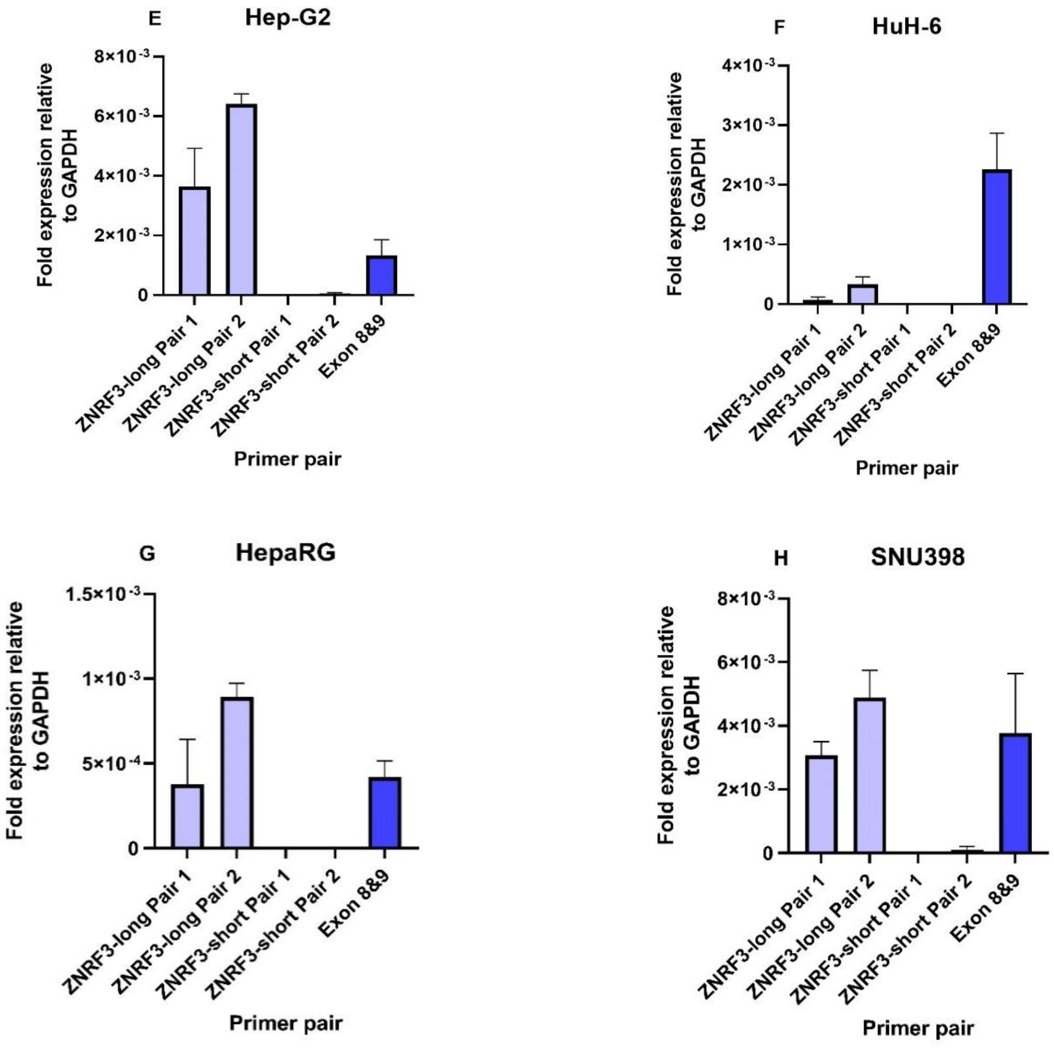

**Supplementary Fig.4 page2**

### Hepatoblastoma/Hepatocellular Carcinoma

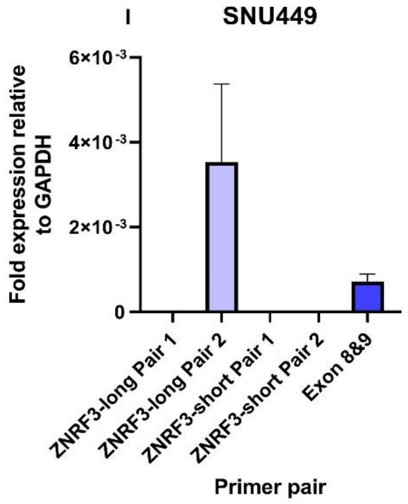


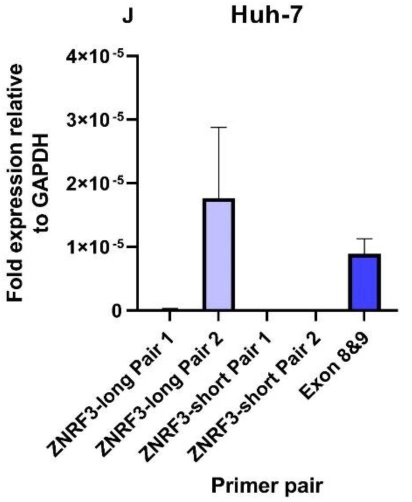


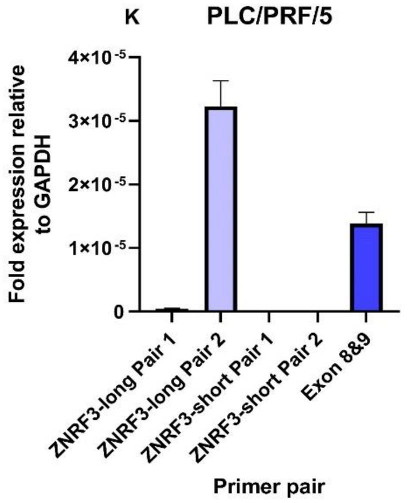


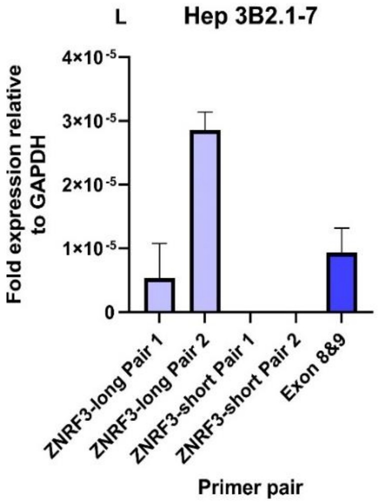


### Pancreatic ductal Carcinoma

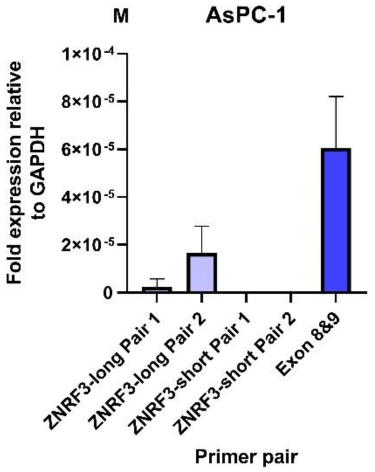


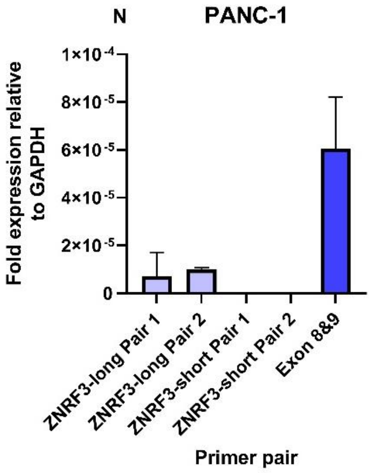


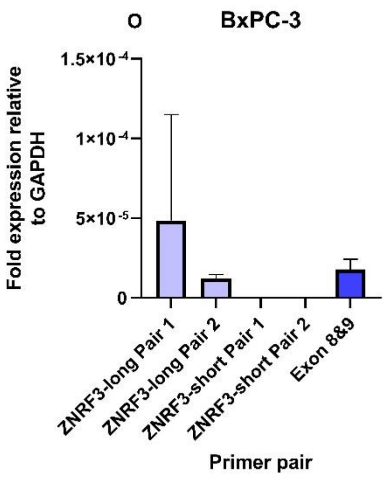


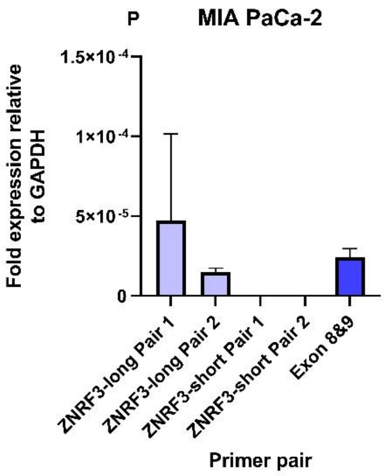

**Supplementary Figure 4**. Comparison of the expression of the different *ZNRF3* RNA isoforms by qPCR. QRT-PCR assay reveals that the long ZNRF3 isoform is constitutively higher expressed compared to the short *ZNRF3* isoform in colon adenocarcinoma (A-D), hepatoblastoma/hepatocellular carcinoma (E-L) and pancreatic ductal carcinoma (M-P) cell lines. Expression levels are relative to the housekeeping gene *GAPDH.* Data are presented as mean ± SD, of three replicates. Two independent primer sets were used for each RNA isoform. Primers based on exon 8&9 used as control.

**Supplementary Fig.5**

#### ZNRF3 long protein isoform

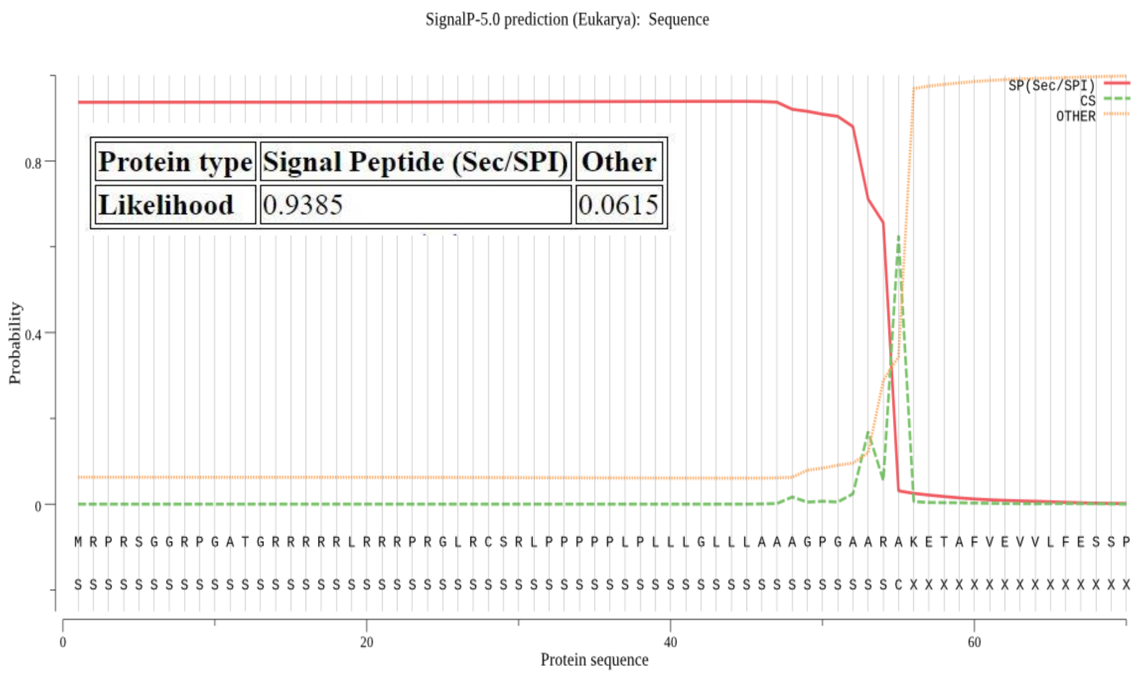


#### ZNRF3 short protein isoform

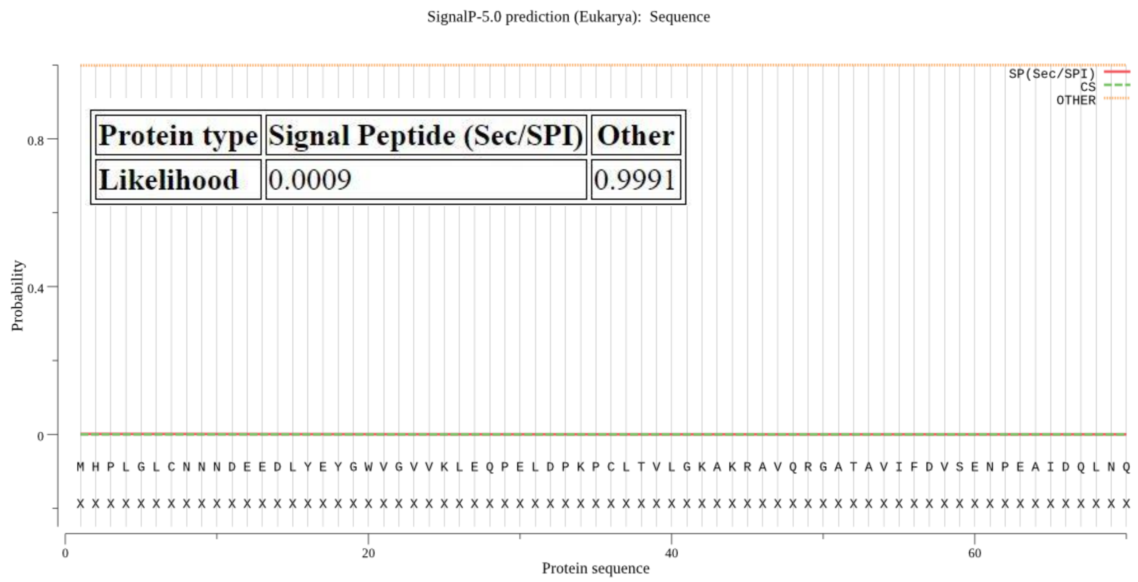


**Supplementary Figure 5**. Prediction of the presence of a signal peptide at the N-terminus of the ZNRF3 protein. For the long isoform, the SignalP-5.0 tool predicts that amino acids 1-55 belong to the signal peptide that is cleaved from the rest of the protein. The lysine (K) residue at position 56 is predicted to be the first amino acid after cleavage of the peptide. The likelihood that this is a signal peptide is 0.939 (max is 1). No signal peptide could be identified in the amino acid sequence of the short ZNRF3 isoform. The likelihood that there is a signal peptide at the N- terminus of this protein is 0.0009. SP: signal peptide; CS: cleavage site.

#### Supplementary Fig.6

**R246*:**

**Clone B8**
CAGCGACGCAGTCAGGTAGTGCCTCTGTGTGTAGCCCGTG (**Parental**)
CAGCGACGCAAGTCAGGTAGTGCCTCTGTGTGTAGCCCGTG (+1bp) p.S247Kfs*55

**Clone B9**
CAGCGACGCAGTCAGGTAGTGCCTCTGTGTGTAGCCCGTG (**Parental**)
CAGCGACGCAAGTCAGGTAGTGCCTCTGTGTGTAGCCCGTG (+1bp) p.S247Kfs*55

**S355*:**

**Clone B2:**
GGAGACCAGCAACCTCTCACGTGGTCGGCAGCAGAGGGTGACCCTGCC (**Parental**)
GGAGACCAGCAACCTCTCTCACGTGGTCGGCAGCAGAGGGTGACCCTGCC (+2bp) p.R356Hfs*7

**Clone C4:**
GGAGACCAGCAACCTCTCACGTGGTCGGCAGCAGAGGGTGACCCTGCC (**Parental**)
GGAGACCAGCA..................gcagagggtgaccctgcc (-19bp) p.N353Sfs*3
GGAGACCAGCA........cgtggtcggcagcagagggtgaccctgcc (-8bp) p.N353Tfs*111
GGAGACCAGCAac.......gtggtcggcaGCAGAGGGTGACCCTGCC (-7bp) p.L354Vfs*6

**S536*:**

**Clone C11:**
TCAGCTGCTATCACGGCCACCGCTCGGTGTGCAGTGGCTACCTGGCCGACTGCCCAGGCAGC (**Parental**)
TCAGCTGCTATCACGGCCACCG.....TGAGCAGTGGCTACCTGGCCGACTGCCCAGGCAGC (-5bp) p.S537Vfs*27
TCAGCTGCTATC............................................GGCAGC (-44bp p.H533Rfs*18

**R789*:**

**Clone A4:**
CTGCTGCTTCTATGAAGAGAAGCAGGTGGCCCGCGGGGGCGGAGGGGGCAGCGG (**Parental**)
CTGCTGCTTCTATGAAGAGAAGCAGGTGGCCCCGCGGGGGCGGAGGGGGCAGCGG (+1bp) p.R789Pfs*11
CTGCTGCTTCTATG.......................GGCGGAGGGGGCAGCGG (-23bp) p.E783Gfs*9

**Clone G2:**
CTGCTGCTTCTATGAAGAGAAGCAGGTGGCCCGCGGGGGCGGAGGGGGCAGCGGCT (**Parental**)
CTGCTGCTTCTATGAAGAGAAG..............GGGCGGAGGGGGCAGCGGCT (-14bp) p.Q786Gfs*9
CTGCTGCTTCTATGAAGAGAAGCAGGTGGCCCGTCGGGGGCGGAGGGGGCAGCGGCT (+1bp) p.G790Rfs*10
CTGCTGCTTCTATGAAGAGAAGCAGGTGGCCCGCCGGGGGCGGAGGGGGCAGCGGCT (+1bp p.G790Rfs*10

**Supplementary Figure 6**. Depiction of CRISPR/Cas9 induced ZNRF3 truncating mutations generated in a RNF43-knockout HEK293T cell line.

#### Supplementary Fig.7

Blast two protein alignment of human RNF43 (783aa) and the long version of ZNRF3 (936aa)

```
RNF43  42   AEQKAIIRVIPLKMDPTGKLNLTLEGV---FAGVAEITPAEGKLMQSHPLYLCNASDDDN  98
            A++ A + V+  +  P+G       G+   F+       AEG+++Q HPL LCN +D+++
ZNRF3  55   AKETAFVEVVLFESSPSGDYTTYTTGLTGRFSRAGATLSAEGEIVQMHPLGLCNNNDEED  114

RNF43  99   L-EPGFISIVKLESPRRAPRPCLSLASKARMAGERGASAVLFDITEDRAAAEQLQQPLG-  156
            L E G++ +VKLE P   P+PCL++  KA+ A +RGA+AV+FD++E+  A +QL Q
ZNRF3  115  LYEYGWVGVVKLEQPELDPKPCLTVLGKAKRAVQRGATAVIFDVSENPEAIDQLNQGSED  174

RNF43  157  -LTWPVVLIWGNDAEKLMEFVYKNQKAHVRIELKEPPAWPD--YDVWILMTVVGTIFVII  213
             L  PVV + G DA KLM  V K + A  RI+ + PP  P   +D+ I +      + ++
ZNRF3  175  PLKRPVVYVKGADAIKLMNIVNKQKVARARIQHR-PPRQPTEYFDMGIFLAFFVVVSLVC  233

RNF43  214  LASVLRIRCRPRHSRPDPLQQRTAWAISQLATRRYQASCRQARGEWPDSGSSCSSAPV--  271
            L  +++I+ + R S+ + + +    A+ ++ TR++ +  +  R        SC +
ZNRF3  234  LILLVKIKLKQRRSQ-NSMNRLAVQALEKMETRKFNSKSKGRRE------GSCGALDTLS  286

RNF43  272  ------CAICLEEFSEGQELRVISCLHEFHRNCVDPWLHQHRTCPLCMFNITEGDSFSQS  325
                  CAICLE++ +G+ELRVI C H FHR CVDPWL QH TCP C  NI E      +
ZNRF3  287  SSSTSDCAICLEKYIDGEELRVIPCTHRFHRKCVDPWLLQHHTCPHCRHNIIEQKGNPSA  346

RNF43  326  LGPSRSYQEPGR--RLHLIRQHPGHAH--YHLPAAYLLGPSRSAVARPPRPGPFLPSQEP  381
            +    S    GR  R+ L   +PG  H    +PA     P+R+++     P   L
ZNRF3  347  VCVETSNLSRGRQQRVTLPVHYPGRVHRTNAIPAY----PTRTSMDSHGNPVTLLTMDRH  402

RNF43  382  GMG---------------PRHHRFPRAAH----------PRAPGEQQRLAG---------  407
            G                 P H     AAH          P  P  + +L+G
ZNRF3  403  GEQSLYSPQTPAYIRSYPPLHLDHSLAAHRCGLEHRAYSPAHPFRRPKLSGRSFSKAACF  462

RNF43  408  ------AQHPYAQGWGLSHLQSTSQH-PAACPVPLRRARPPDSSGS--------------  446
                   QH Y Q  GLS+ +   Q  P+  P    RA PP  SGS
ZNRF3  463  SQYETMYQHYYFQ--GLSYPEQEGQSPPSLAPRGPARAFPPSGSGSLLFPTVVHVAPPSH  520

RNF43  447  GESYCTER-----------SGYLADGP----ASDSSSGPCHGSSSDSVVNCTDISLQGVH  491
             ES  T             SGYLAD P    +S SSSG CH SSSDSVV+CT++S QGV+
ZNRF3  521  LESGSTSSFSCYHGHRSVCSGYLADCPGSDSSSSSSSGQCHCSSSDSVVDCTEVSNQGVY  580

RNF43  492  GSSSTFCSSLSSDFDPLVY  510
            GS STF SSLSSD+DP +Y
ZNRF3  581  GSCSTFRSSLSSDYDPFIY  599
```

- Variants behaving as WT
- LOF(or partial LOF) and Hyper-activating mutations
- R-spondin binding domain
- RING domain

**Supplementary Figure 7**. Human RNF43-ZNRF3 protein alignment. The Rspondin binding and RING domains are the most conserved between RNF43 and ZNRF3, in addition to the S-rich domain. 17 of 51 mutations in the RNF43 Rspondin binding domain and 12 of 47 mutations in the ZNRF3 Rspondin binding domain are (partial) loss-of-function or hyper-activating mutations. For the RING domain of RNF43, 39 of 44 variants behave as loss-of-function or hyper-activating mutations, while this is the case for 14 of 27 ZNRF3 Ring domain variants.

**Supplementary Fig.8**

### ConSurf Results based on human ZNRF3

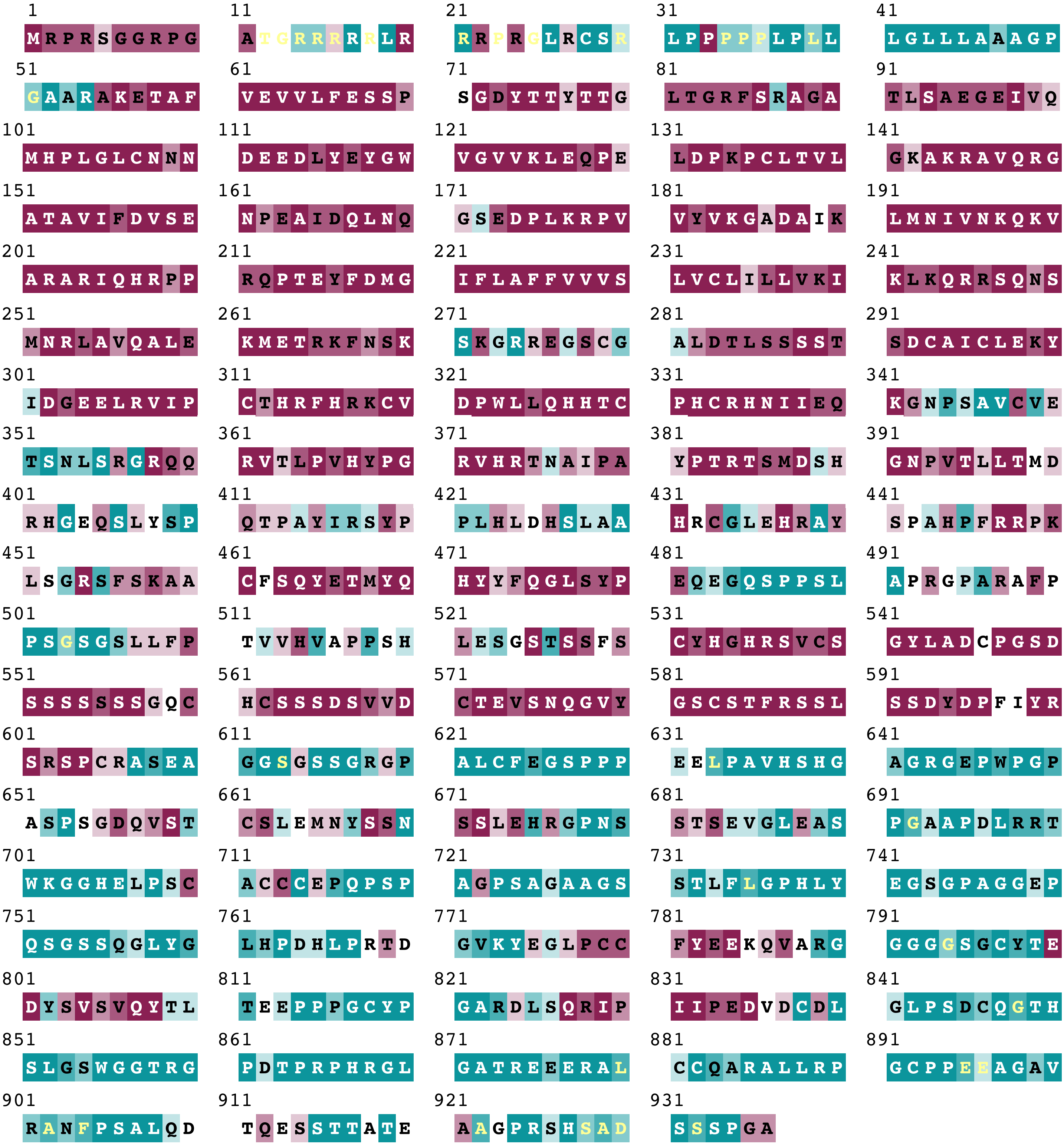


**The conservation scale:**

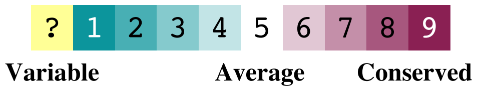


**Supplementary Figure 8.** Analysis of evolutionary conserved ZNRF3 residues using the ConSurf tool. Analysis is based on the human 936 amino acid isoform of ZNRF3 (Uniprot Q9ULT6-1).

#### Supplementary Fig.9

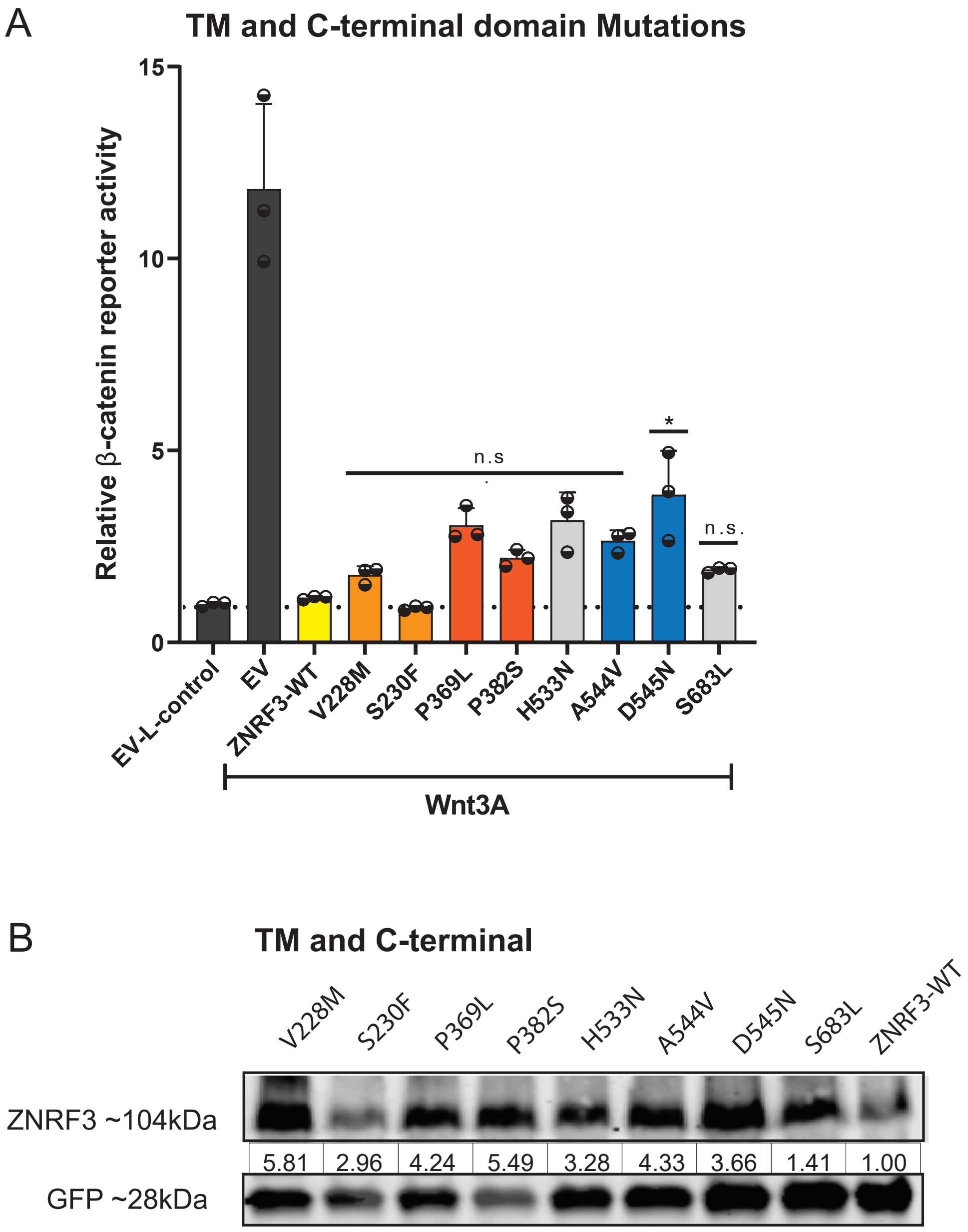


**Supplementary Figure 9.** All tested variants in the ZNRF3 transmembrane (TM) and C-terminal domains behave similar to wild-type ZNRF3. **(A)** A β-catenin reporter assay reveals that 7/8 variants are comparable in Wnt/β-catenin signaling as wild-type. Only the D545N shows a significant albeit modest defect in functionality. Wnt3A conditioned medium was added to Empty Vector (EV) and all ZNRF3 variant transfected cells. L-control medium was added to EV transfected cells, serving as a negative control for β-catenin signaling. All relative β-catenin reporter activities are depicted as WRE/CMV-Renilla ratios, in which the value obtained for ZNRF3-WT was arbitrarily set to 1 (one-way ANOVA; * $P<0.1$). **(B)** All variants were co-transfected with EGFP plasmids. In the immunoblot, all variants show increased protein stability. All relative protein stabilities are depicted as ZNRF3 variant/GFP ratios, in which the value obtained from ZNRF3-WT was set to 1.

#### Supplementary Fig.10

A

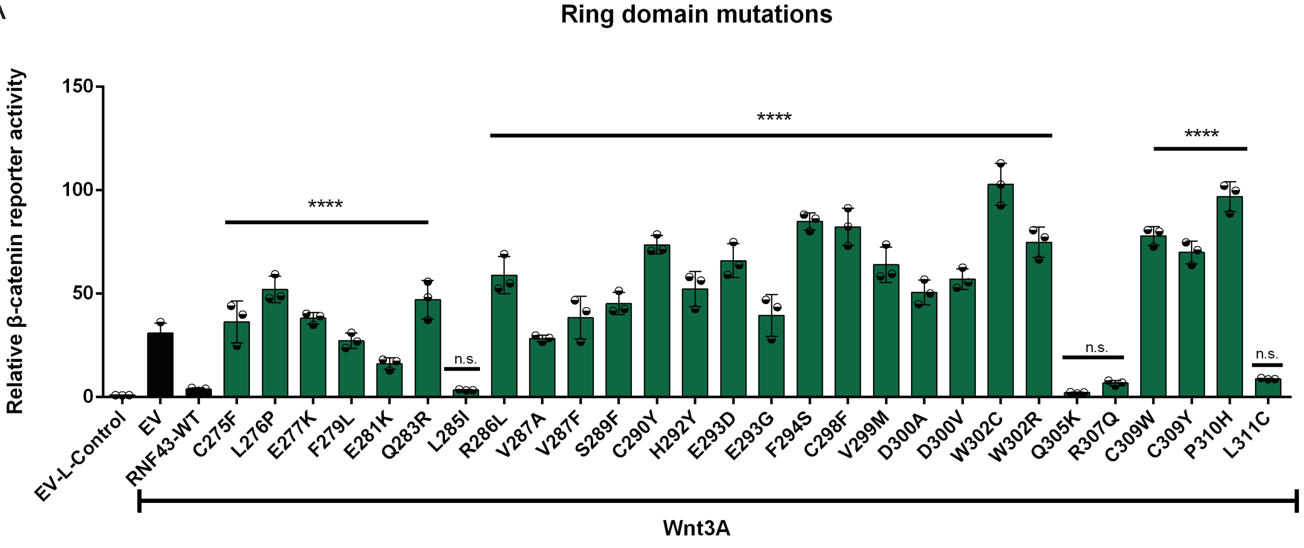


B

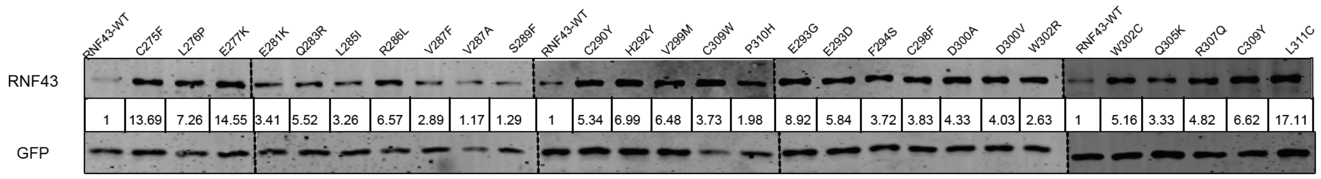


**Supplementary Figure 10.** Analysis of 28 previously unexplored tumor-associated RING domain variants for RNF43. **(A)** A β-catenin reporter assay reveals that 24 out of 28 variants behave as loss-of-function or hyperactivating mutants. Wnt3A conditioned medium was added to Empty Vector (EV) and all RNF43 variant transfected cells. L-control medium was added to EV transfected cells as a negative control for β-catenin signaling. All relative β-catenin reporter activities are depicted as WRE/CMV-Renilla ratios, in which the value obtained for RNF43-WT was arbitrarily set to 1. Statistical significance was assessed using a one-way ANOVA test (****$P<0.0001$). **(B)** All variants were co-transfected with EGFP plasmids. In the immunoblot, basically all variants show increased protein stability. All relative protein stabilities are depicted as RNF43/GFP ratios, in which the value obtained from RNF43-WT was set to 1.

#### Supplementary Fig.11

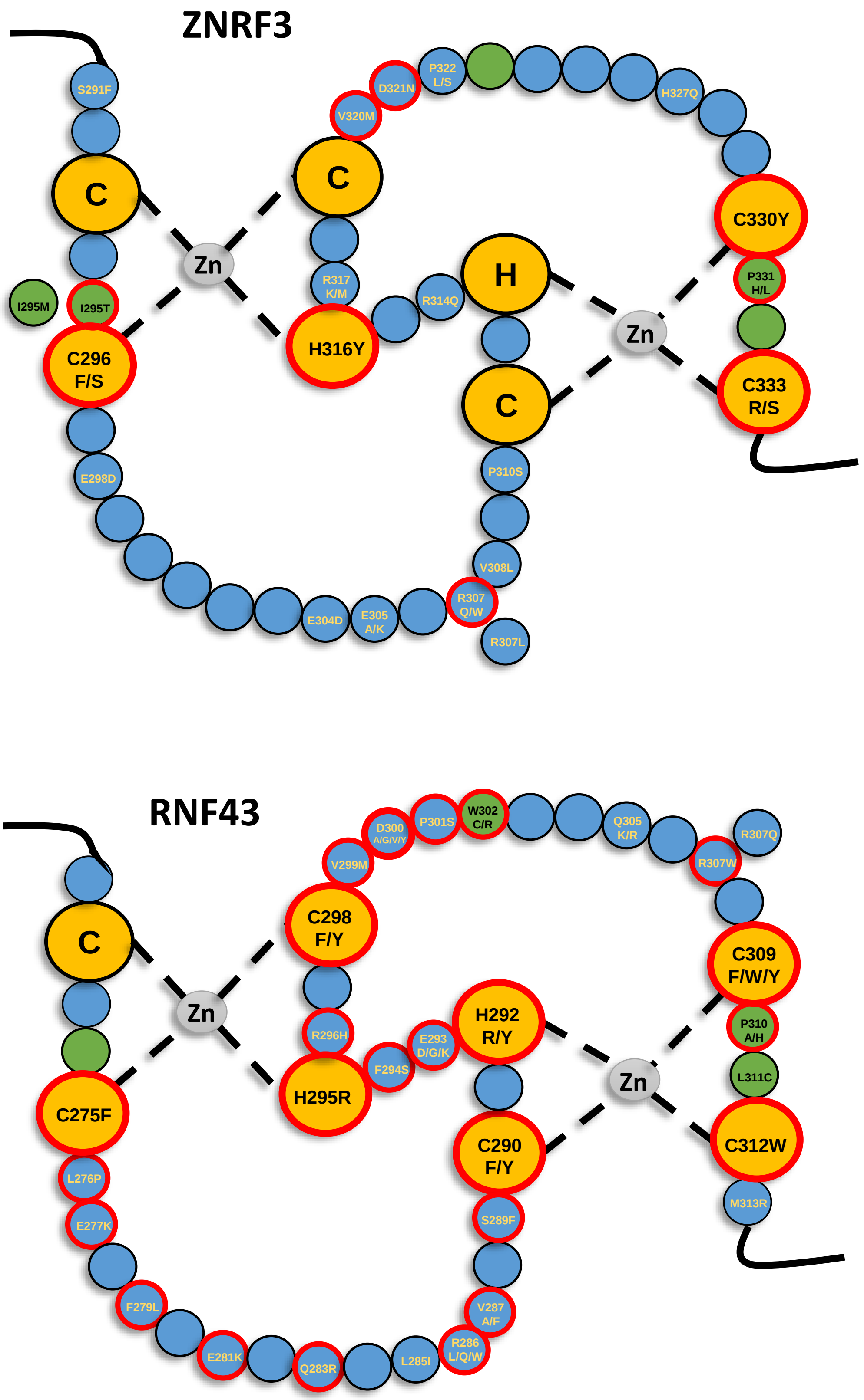


**Supplementary Figure 11**. Schematic representation of the RING domain structures of ZNRF3 and RNF43. The green circles are the residues required for interaction with ubiquitin-conjugating E2 enzymes, while the yellow circles are the C/H residues involved in Zinc (Zn) coordination. Tumor-associated variants that disrupt the β-catenin regulatory function of ZNRF3/RNF43 are encircled by a red border.

### Supplementary Fig.12

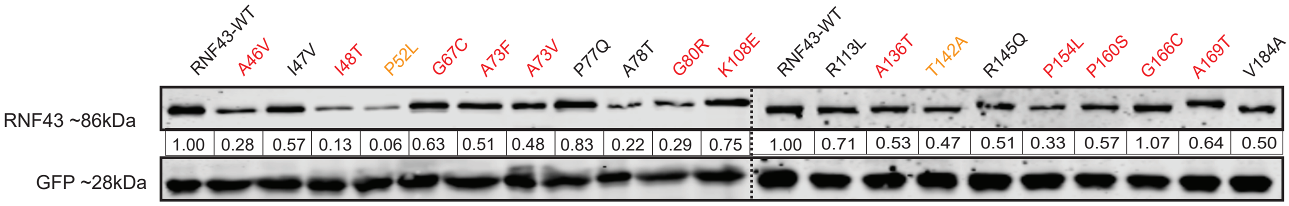


**Supplementary Figure 12.** Analysis of protein stability of 20 tumor-associated Rspondin binding domain variants for RNF43. All variants were co-transfected with EGFP plasmid. Variants with similar functionality as wild-type are depicted in black font, while LOF/hyperactivating or partially defective variants are depicted in red or orange font, respectively. Functionality is based on results previously published by Yu et al.[1]. In the immunoblot, several defective variants show reduced protein stability, although exceptions are observed (e.g. G166C), and also some variants behaving as wild-type show a weak or stronger (e.g. A78T) reduction in protein levels. Overall a trend is observed of reduced stability of defective variants, although not consistently. All relative protein stabilities are depicted as RNF43/GFP ratios, in which the value obtained from RNF43-WT was set to 1.


1 *Yu, J. et al.* The Functional Landscape of Patient-Derived RNF43 Mutations Predicts Sensitivity to Wnt Inhibition. Cancer Res **80**, 5619-5632, doi:10.1158/0008-5472.Can-20-0957 (2020).

#### Supplementary Fig.13

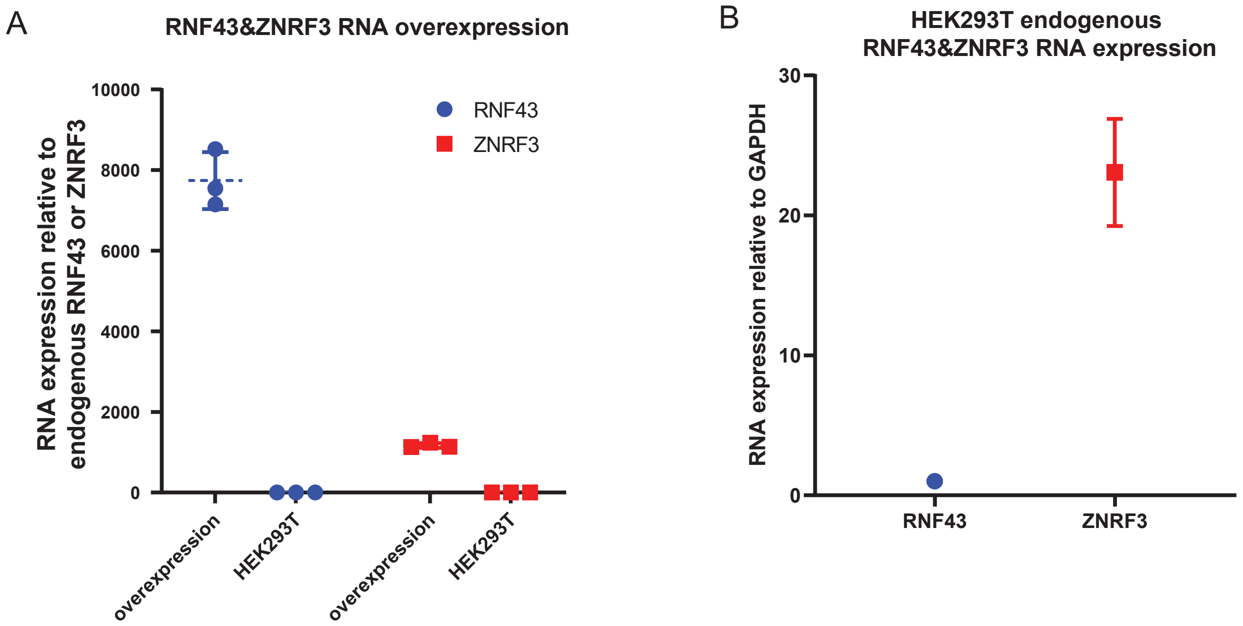


**Supplementary Figure 13.** Relative mRNA levels of *RNF43* and *ZNRF3* in HEK293T cells. **(A)** Our transient transfection experiments lead to an approximately 8000- and 1600-fold higher expression of *RNF43* and *ZNRF3*, respectively. **(B)** At the endogenous level, *ZNRF3* expression is about 20-30-fold more than *RNF43* in HEK293T cells (in triplicate, n=3, normalized to *GAPDH*).

#### Supplementary Fig.14

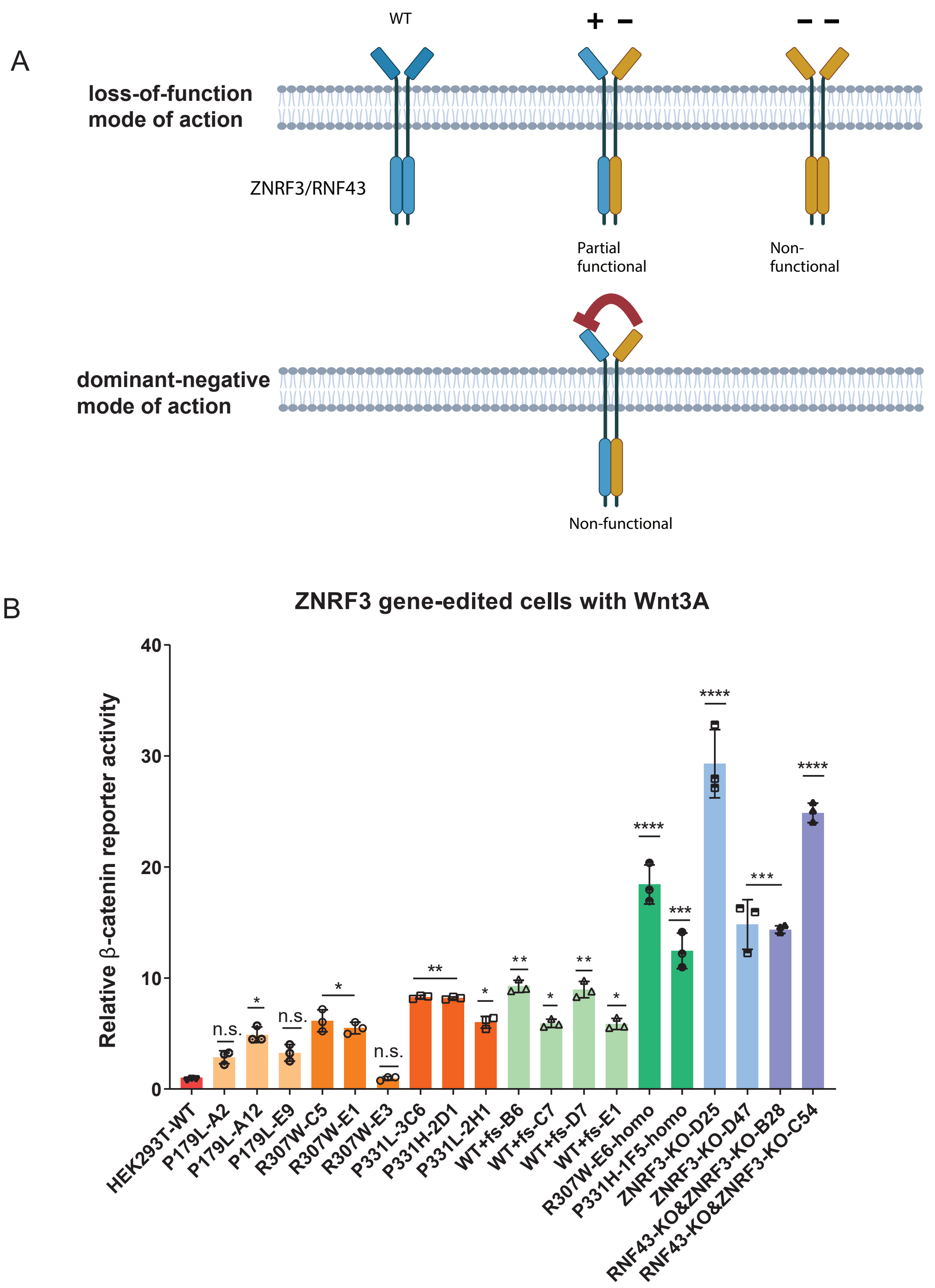


**Supplementary Figure 14.** Endogenous levels of heterozygous missense mutations do not affect ZNRF3 function more strongly than heterozygous knockout mutations. **(A)** ZNRF3 and RNF43 have been shown to form hetero- and homodimers. Theoretically, two scenarios can be envisaged leading to loss of RNF43/ZNRF3 functionality based on this dimer formation. First, when a dimer is formed between a wild-type and mutant RNF43/ZNRF3, the dimer will only lose half its function. Only when both proteins are mutant, it will lead to a non-functional dimer. Second, the presence of a mutant RNF43/ZNRF3 in the dimer leads to a dominant-negative inhibition of the entire dimer. This latter scenario has been considered to be the most likely based on the observed hyperactivation of β-catenin signaling when a mutant variant is overexpressed. **(B)** β-catenin reporter assays for all individual knockin/out ZNRF3 clones. All heterozygous mutations, including P179L, R307W, P331L or P331H, and WT+frameshift (fs), slightly increase β-catenin signaling (2-8 fold) to similar levels. There is one outlier clone, i.e. R307W-E3, which shows no clear increase in signaling. Homozygous ZNRF3 knockout clones (D25, D47), homozygous R307W or P331H knockin clones (E6, 1F5), and RNF43/ZNRF3 double knockout clones, all show a comparable further increase in signaling (18-30 fold). The relative Wnt/β-catenin signaling activities are depicted as WRE/CMV-Renilla ratios, in which the value obtained for wild-type (WT) HEK293T cells was arbitrarily set to 1. Wnt3A conditioned medium was added to WT HEK293T cells and all CRISPR-edited HEK293T cells. Statistical significance was assessed using a one-way ANOVA test (****P<0.0001, ***P<0.001, **P<0.01, *P<0.1).

**Supplementary Fig.15**

**Original IB for Fig.2**

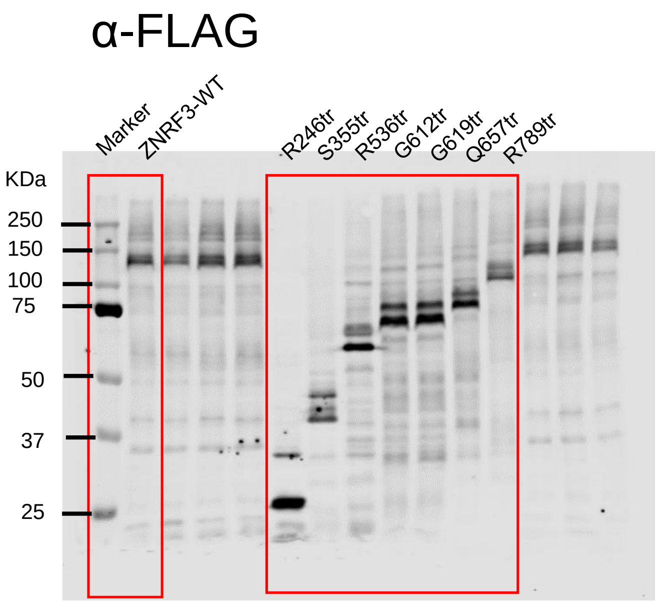

**Supplementary Fig.15, continued**

**Original IB for Fig.3**

HA-tag

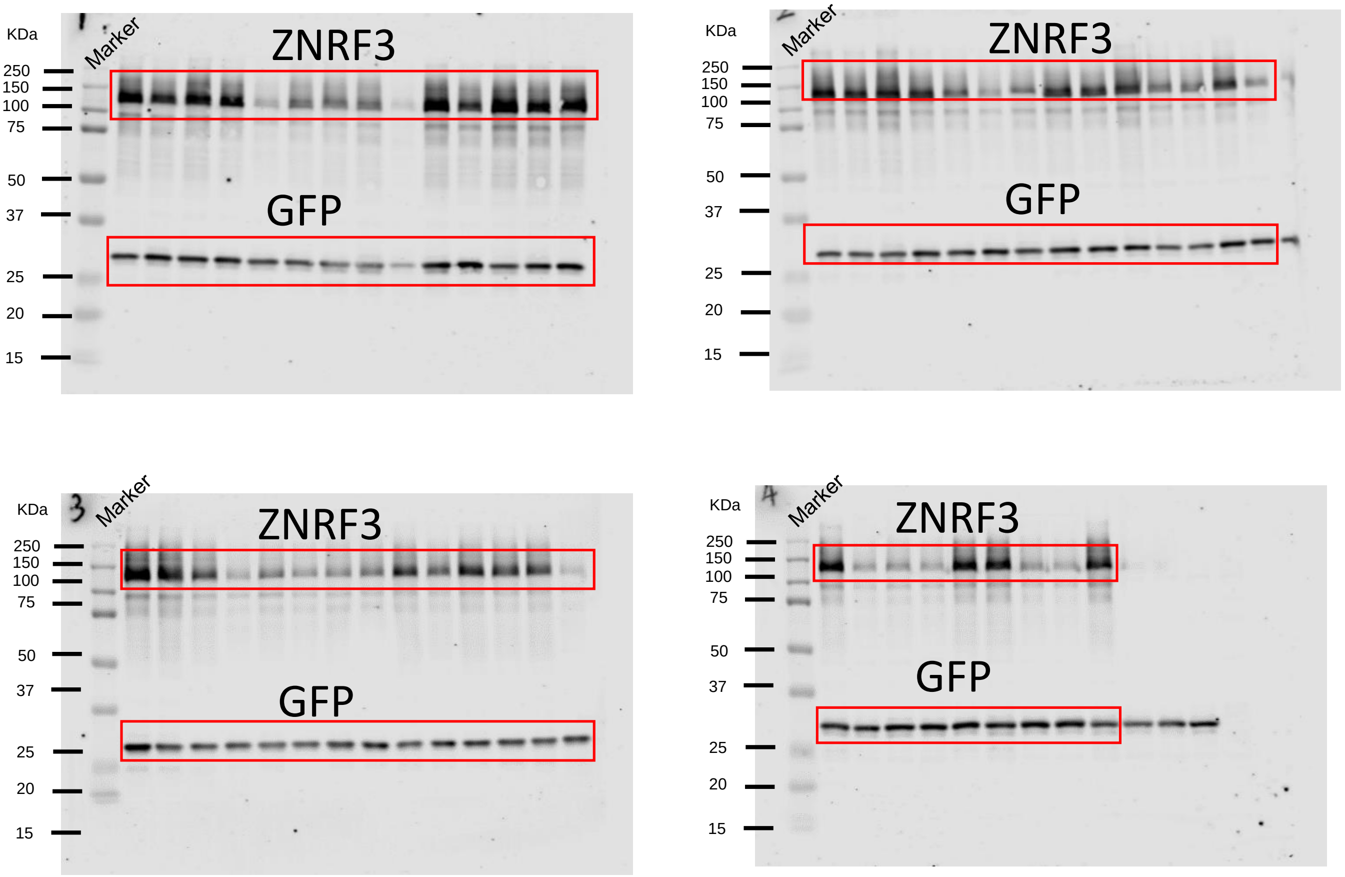

**Supplementary Fig.15, continued**

**Original IB for Fig.4**

α-FLAG

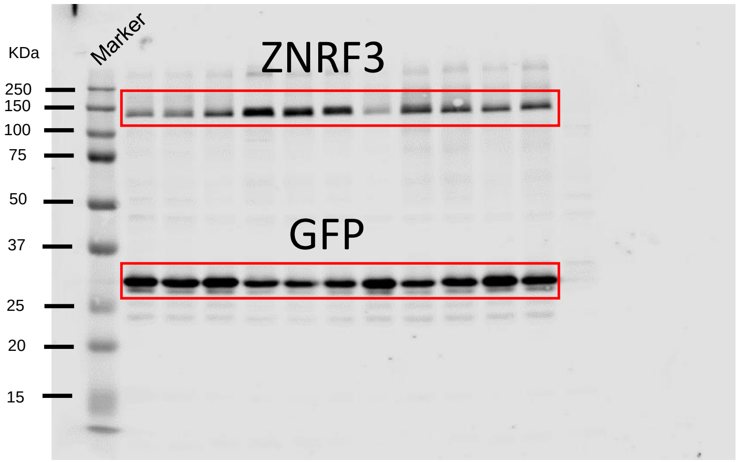


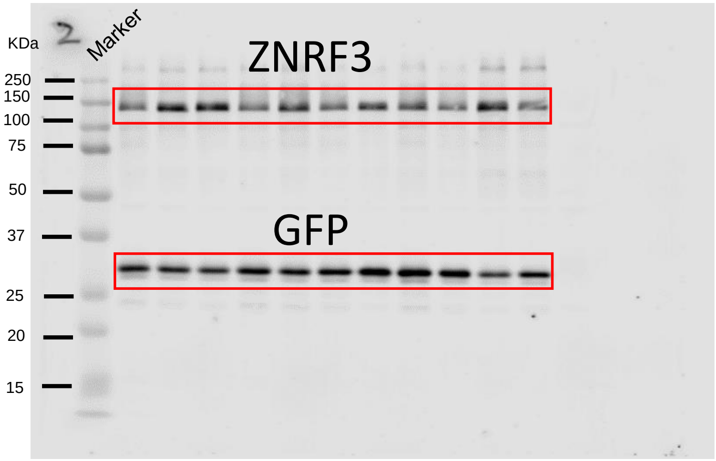


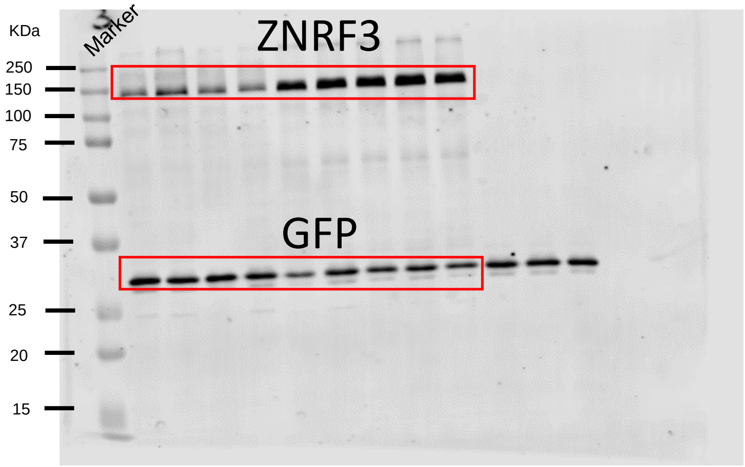

**Supplementary Fig.15, continued**

**Original IB for Fig.5**

#### IP: Streptavidin

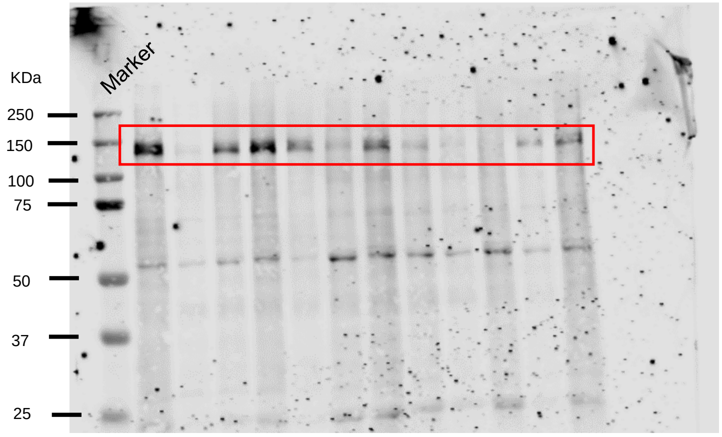


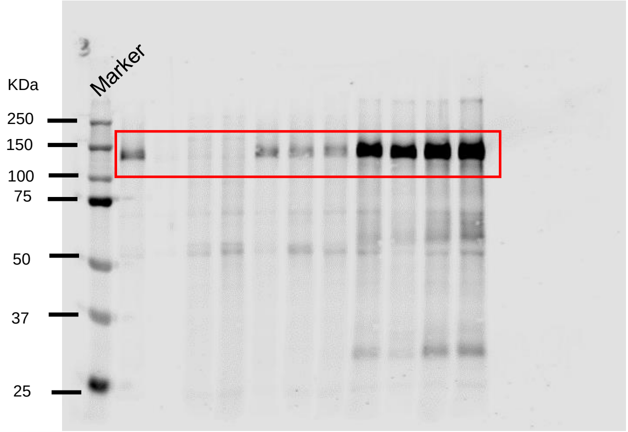


#### IP: α-FLAG

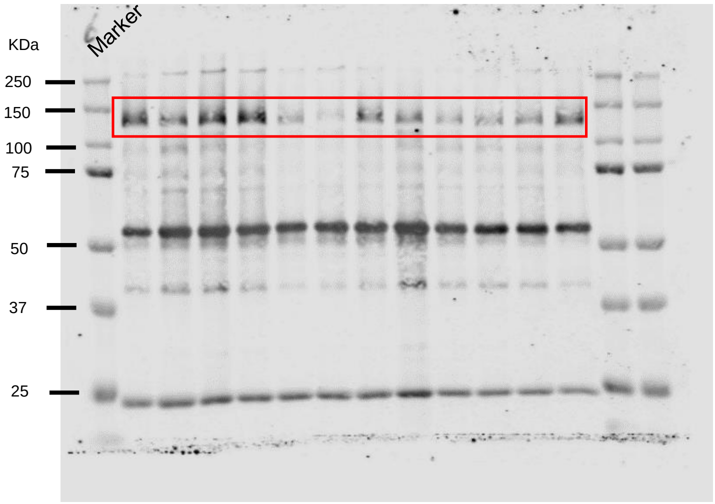


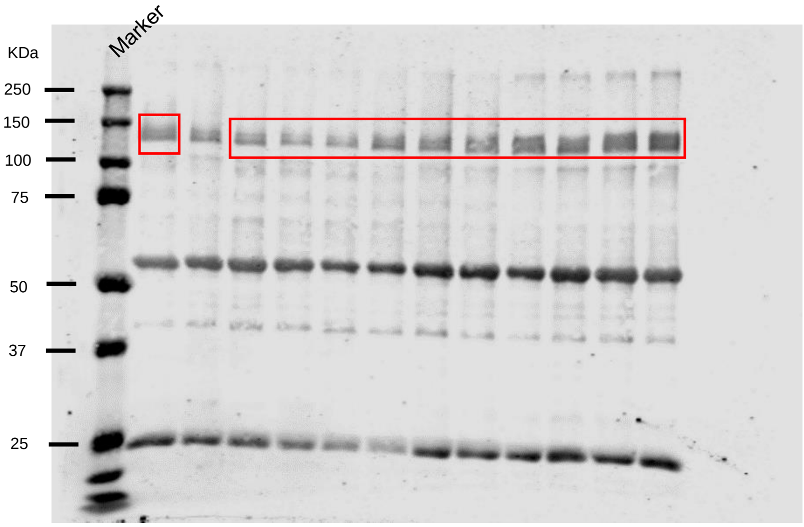

**Supplementary Fig.15, continued**

**Original IB for Fig.6**

#### a. HEK293T cells at 37°C

IP: α-FLAG

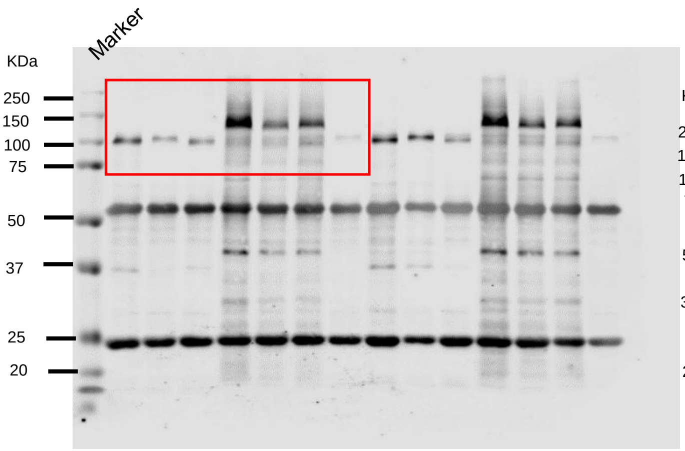


IP: Streptavidin

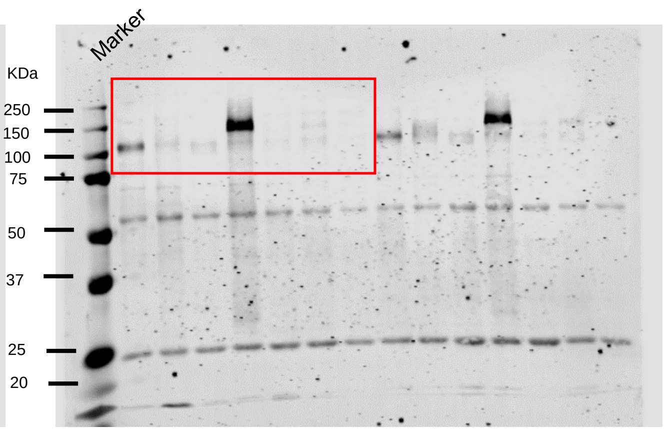


Input: α-FLAG

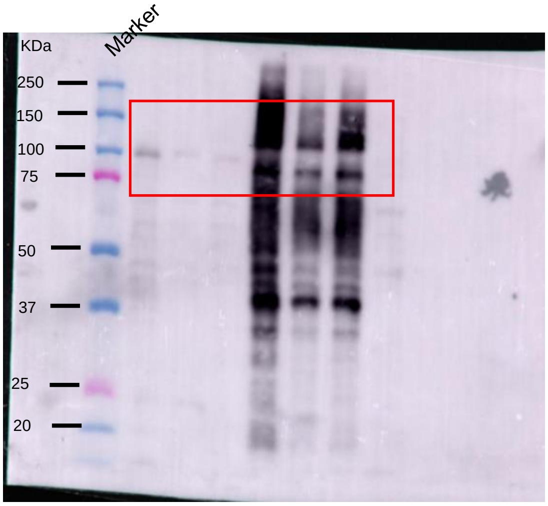


Input: Streptavidin

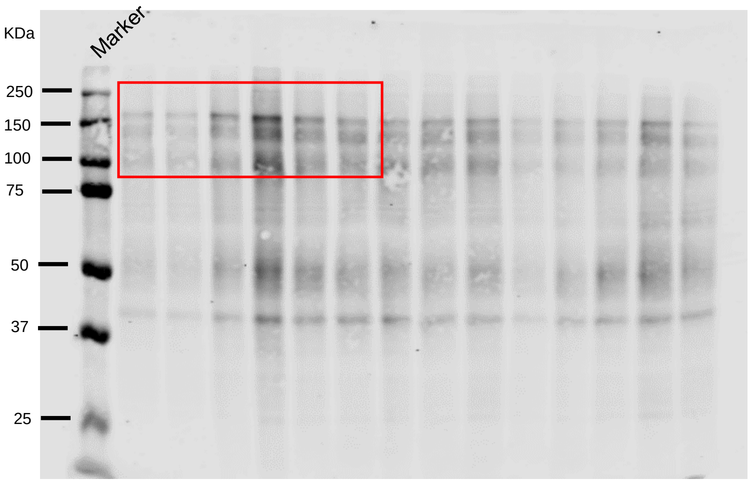

**Supplementary Fig.15, continued**

**Original IB for Fig.6**

a. HEK293T cells at 27°C

IP: α-FLAG

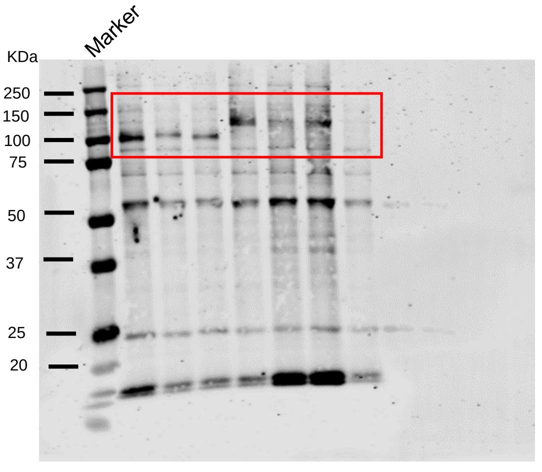


IP: Streptavidin

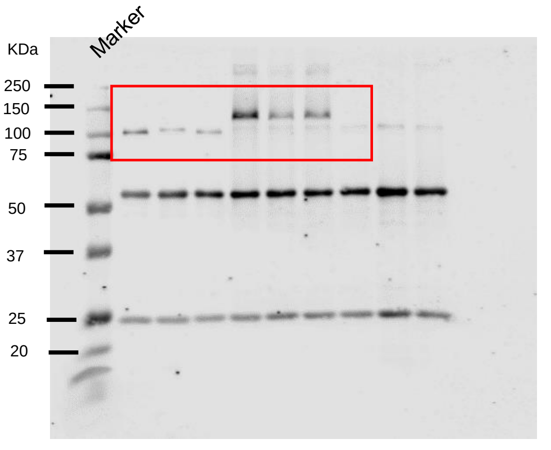


Input: α-FLAG

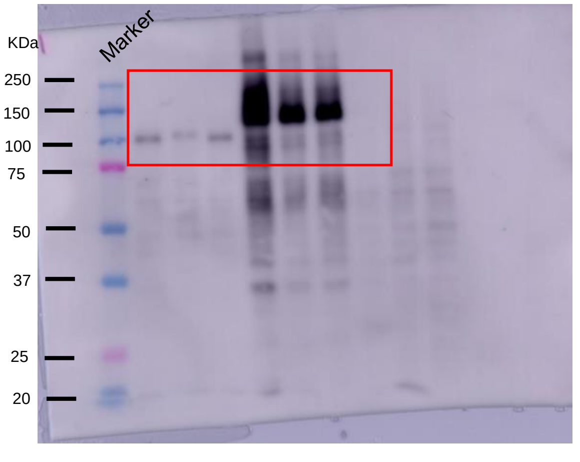


Input: Streptavidin

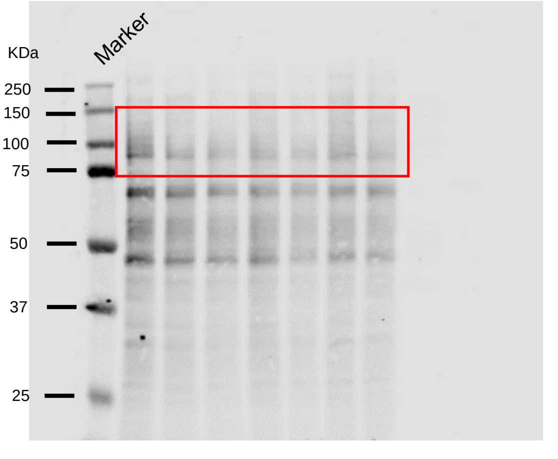

**Supplementary Fig.15, continued**

**Original IB for Supplementary Fig.9**

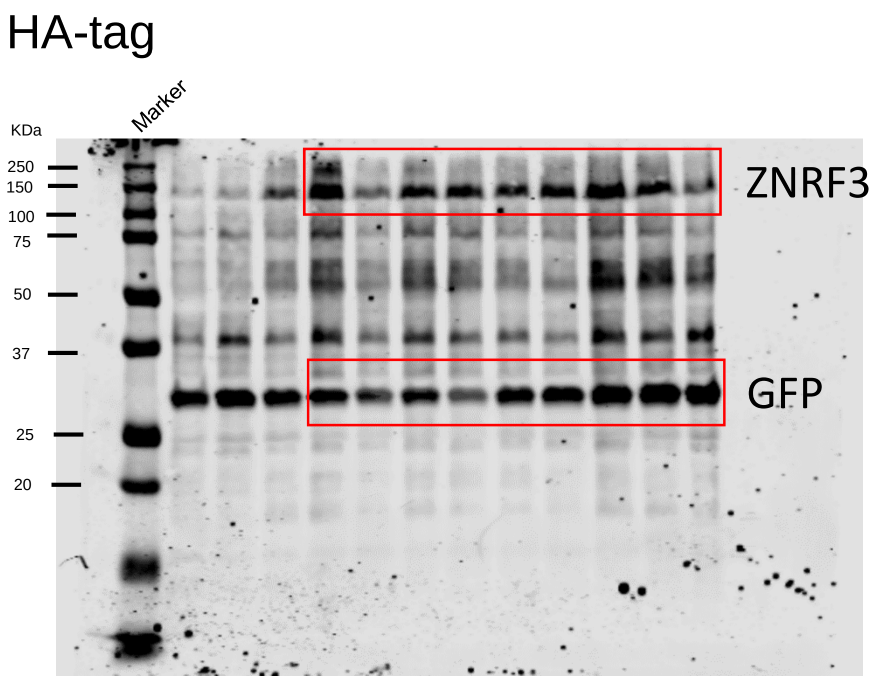

**Supplementary Fig.15, continued**

**Original IB for Supplementary Fig.10**

α-FLAG

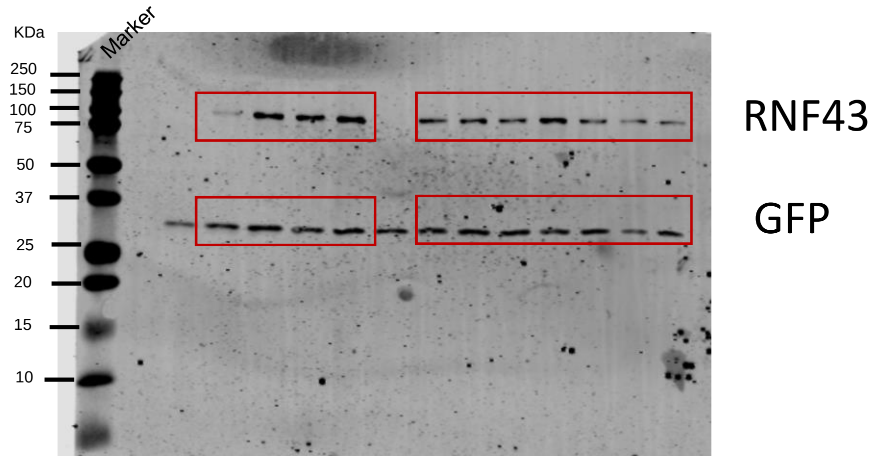


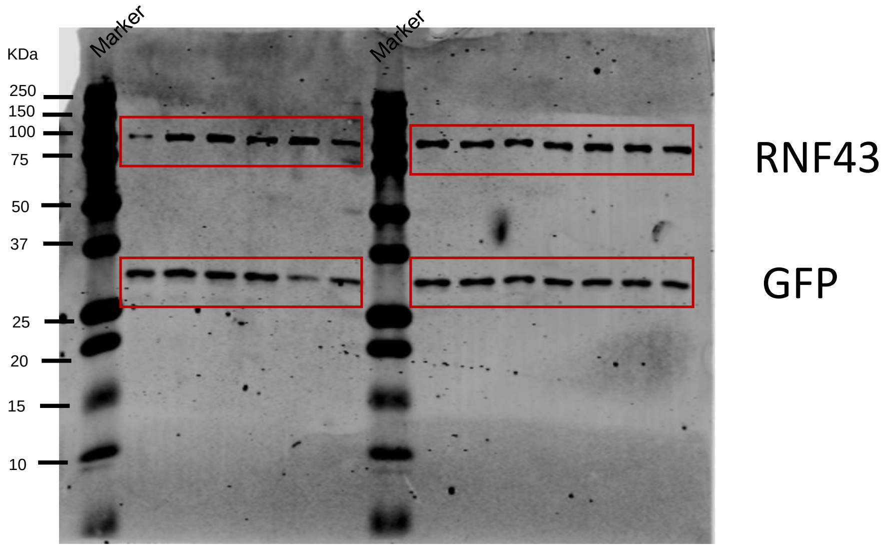


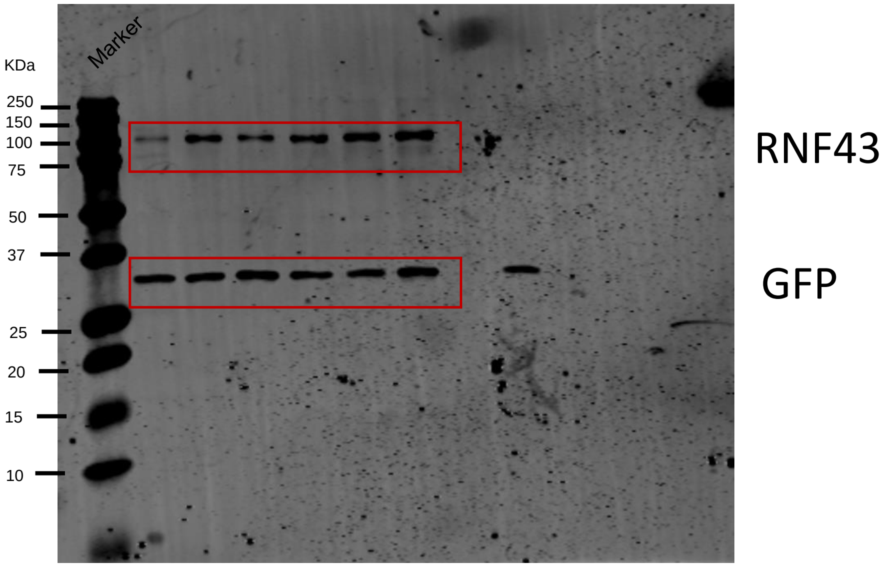

**Supplementary Fig.15, continued**

**Original IB for Supplementary Fig.12**

α-FLAG

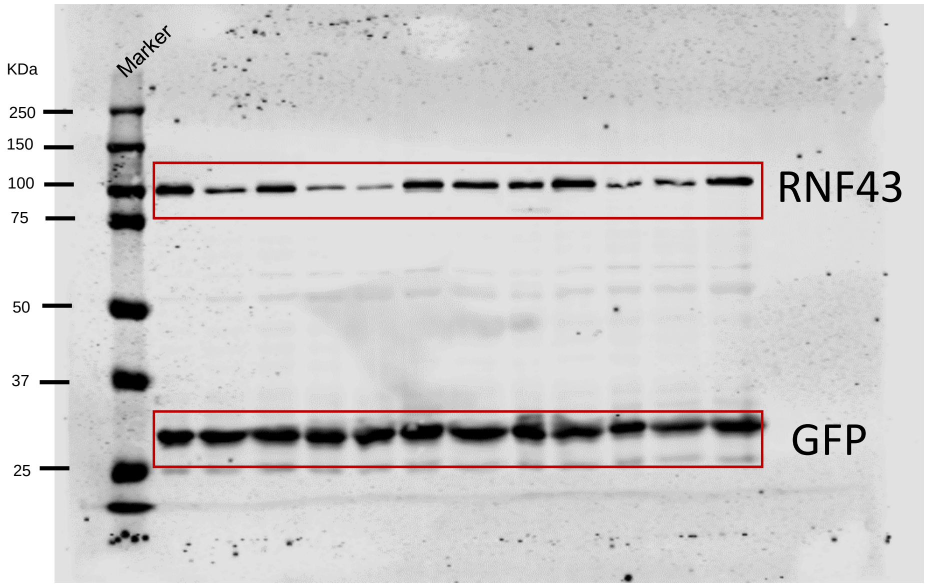


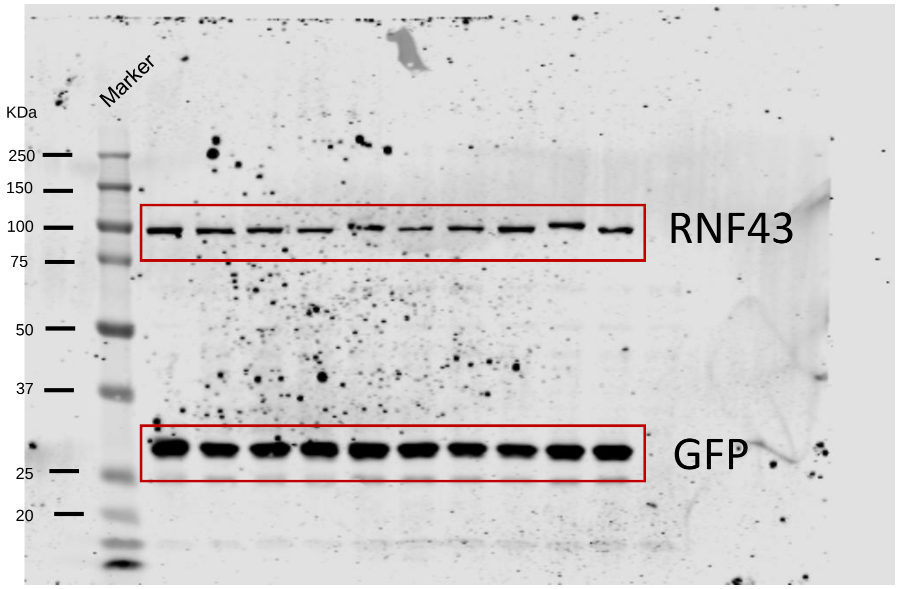
