## Supplemental Tables for "Lack of dominant-negative activity for tumor-associated ZNRF3 missense mutations at endogenous expression levels"

### Supplementary Table 1

| domain | Cancer type | ZNRF3 AA change | Conservation (1-9) | Function |
|---|---|---|---|---|
| R-spondin binding | Lung Carcinoma | E67Q | 9 | |
| | Pancreatic Adenocarcinoma | S68W | | |
| | Upper aerodigestive trac Carcinoma | S69N | | |
| | | G83D | 8 | LOF |
| | Thyroid Carcinoma | A88V | | |
| | Biliary tract Carcinoma | A94T | | |
| | Skin Carcinoma | E95K | | |
| | Lung Squamous Cell Carcinoma | E95Q | | |
| | Uterine Endometrioid Carcinoma | M101I | 9 | |
| | | M101T | | |
| | | P103A | | |
| | Lung Adenocarcinoma | G105C | | LOF |
| | Renal Clear Cell Carcinoma | G105D | | |
| | Uterine Endometrioid Carcinoma /Colorectal Adenocarcinoma | E112K | | |
| | Glioblastoma | E113K | | |
| | Biliary tract Carcinoma | E117K | | |
| | Skin Carcinoma | W120L | | |
| | Thyroid/ Urinary tract Carcinoma | V121G | | LOF |
| | | K125E | | |
| | Cutaneous Melanoma | D132N | | |
| | Cutaneous Melanoma | P135Q | | |
| | Adrenal cortical carcinoma | G141S | | |
| | Colon Adenocarcinoma | A143T | | |
| | Uterine Serous Carcinoma/Uterine Papillary Serous Carcinoma | A143V | | |
| | Liver Neoplasm | K144E | | |
| | Hepatocellular Carcinoma | R149P | | LOF |
| | Pancreas Insulinoma | R149Q | | |
| | Uterine Endometrioid Carcinoma | R149W | | |
| | Cutaneous Melanoma | G150V | | LOF |
| | Hepatocellular Carcinoma | A151D | | LOF |
| | | A151P | | |
| | | A153S | | |
| | | A153V | | |
| | Lung Adenocarcinoma | E160Q | | |
| | Colorectal Adenocarcinoma | N169S | | |
| | Stomach Carcinoma | D174E | | |
| | Biliary tract Carcinoma | P175L | 9 | |

| | | | | |
|---|---|---|---|---|
| | Cutaneous Melanoma/Hepatocellular Carcinoma | P175Q | | |
| | Colon Adenocarcinoma | P179L | | Hyper-activiting |
| | Breast Carcinoma | V180G | | LOF |
| | Large intestine Carcinoma | A188D | | LOF |
| | Colon Adenocarcinoma& Esophagogastric Adenocarcinoma | A188V | | LOF |
| | Endometrial Carcinoma | K197T | | |
| | Hepatocellular Carcinoma | A201S | | |
| | Large intestine Carcinoma | R202P | | LOF |
| | Colorectal Adenocarcinoma | I205T | | LOF |
| | Intrahepatic Cholangiocarcinoma | D218G | | |
| TM | Breast Invasive Ductal Carcinoma | V228M | | |
| | Angiosarcoma | S230F | | |
| RING | Desmoplastic Melanoma | S291F | 8 | |
| | Parathyroid Carcinoma | I295M | | |
| | Uterine Serous Carcinoma/Uterine Papillary Serous Carcinoma | I295T | 9 | Hyper-activiting |
| | Large intestine Carcinoma | C296F | | |
| | Lung Adenocarcinoma | C296S | | |
| | Breast Carcinoma | E298D | | |
| | Uterine Endometrioid Carcinoma | E304D | | |
| | Large intestine Carcinoma | E305A | | |
| | Head and Neck Squamous Cell Carcinoma | E305K | | |
| | Large intestine Carcinoma | R307L | | |
| | Colorectal Adenocarcinoma/ Colon Adenocarcinoma | R307Q | | LOF |
| | Adrenocortical Carcinoma/ Uterine Endometrioid Carcinoma/ Uterine Serous Carcinoma/Uterine Papillary Serous Carcinoma/ Intestinal Type Stomach Adenocarcinoma | R307W | 9 | LOF |
| | Pleural Mesothelioma | V308L | 9 | |
| | Cutaneous Melanoma | P310S | | |
| | Stomach Adenocarcinoma | R314Q | | |
| | Esophageal Squamous Cell Carcinoma | H316Y | | Hyper-activiting |
| | Large intestine Carcinoma | R317K | | |
| | Uterine Serous Carcinoma/Uterine Papillary Serous Carcinoma | R317M | 8 | |
| | Colorectal Adenocarcinoma/ Stomach Adenocarcinoma | V320M | 9 | LOF |
| | Uterine Endometrioid Carcinoma | D321N | | LOF |

| | | | | |
|---|---|---|---|---|
| | Skin Malignant melanoma | P322L | | |
| | Cutaneous Melanoma | P322S | | |
| | Large intestine Carcinoma | H327Q | | |
| | Prostate Adenocarcinoma/ Uterine Endometrioid Carcinoma | C330Y | | Hyper-activiting |
| | Colorectal Adenocarcinoma | P331H | | |
| | Skin Cancer, Non-Melanoma | P331L | | |
| | Stomach Adenocarcinoma | C333R | | |
| | Hepatocellular Carcinoma | C333S | | |
| DIR | Skin Cancer, Non-Melanoma | P369L | | |
| | Melanoma /Colorectal Adenocarcinoma/ Melanoma of Unknown Primary | P382S | | |
| Between DIR and S-rich | Intrahepatic Cholangiocarcinoma | H533N | | |
| S-rich | Melanoma | A544V | | |
| | Bladder Urothelial Carcinoma | D545N | | |
| C-teminus | Lung Squamous Cell Carcinoma/ Bladder Urothelial Carcinoma | S683L | | |

### Supplementary Table 2. Primers used to introduce truncating mutations in the *ZNRF3* gene

| | |
|---|---|
| ZNRF3-FLAGtr-Forward | TCCGGAGGTGGATCGGGAG |
| ZNRF3-R246tr-Reverse | CTGCTTCAGCTTGATTTTGACAAGG |
| ZNRF3-S355tr-Reverse | TGAGAGGTTGCTGGTCTCC |
| ZNRF3-R536tr-Reverse | GTGGCCGTGATAGCAGCTG |
| ZNRF3-G612tr-Reverse | CCCCGCCTCACTGGCACG |
| ZNRF3-G619tr-Reverse | CCGGCCCGAGCTGCCCGA |
| ZNRF3-Q657tr-Reverse | ATCCCCCGAGGGAGAGGC |
| ZNRF3-R789tr-Reverse | GGCCACCTGCTTCTCTTCATAG |

### Supplementary Table 3. Primers used to introduce missense mutations in the *ZNRF3* gene

The table below shows the primers that were used to introduce the mutations into the *ZNRF3* coding sequence. The changed codons are shown in red.

| Mutation | Domain | Forward primer | Reverse primer | Tm |
|---|---|---|---|---|
| G83D | R-spondin binding | GGCCTCACGgacCGCTTCTCG | GGTGGTGTAGGTGGTGTAATCG | 69°C |
| E95K | R-spondin binding | GCTCAGCGCCaagGGCGAGATCGTGC | GTGGCCCCGGCCCGCGAG | 72°C |

| | | | | |
|---|---|---|---|---|
| E95Q | R-spondin binding | GCTCAGCGCCcagGGCGAGATCGTGC | GTGGCCCCGGCCCGCGAG | 72°C |
| P103A | R-spondin binding | GCAGATGCACgcaCTGGGCCTATG | ACGATCTCGCCCTCGGCG | 72°C |
| K125E | R-spondin binding | AGGAGTGGTGgagCTGGAACA | ACCCAGCCATATTCATACAAG | 66°C |
| P135Q | R-spondin binding | GGACCCGAAAcaaTGCCTCACTG | AATTCTGGCTGTTCCAGCTTC | 66°C |
| A143T | R-spondin binding | CCTAGGCAAGaccAAGCGAGCAG | ACAGTGAGGCATGGTTTCG | 66°C |
| R149P | R-spondin binding | AGCAGTACAGccgGGAGCTACTG | CGCTTGGCCTTGCCTAGG | 67°C |
| G150V | R-spondin binding | AGTACAGCGGgtaGCTACTGCAG | GCTCGCTTGGCCTTGCCT | 69°C |
| A153S | R-spondin binding | GGGAGCTACTtcaGTCATCTTTGATG | CGCTGTACTGCTCGCTTG | 65°C |
| E160Q | R-spondin binding | TGATGTGTCTcaaAACCCAGAAG | AAGATGACTGCAGTAGCTC | 61°C |
| P179L | R-spondin binding | GCTCAAGAGGctgGTGGTGTATG | GGGTCTTCAGAGCCCTGG | 69°C |
| A188V | R-spondin binding | GGGTGCAGATgtcATTAAGCTGA | TTCACATACACCACCGGC | 66°C |
| I205T | R-spondin binding | TCGAGCAAGGaccCAGCACCGCC | GCCACTTTCTGCTTGTTGACGATGTT | 72°C |
| D218G | R-spondin binding | TGAATACTTTggcATGGGGATTTTCC | GTGGGTTGTCGAGGAGGG | 65°C |
| S291F | RING | AGCTCCACGttcGACTGTGCC | GCTGCTGAGTGTGTCCAGG | 69°C |
| I295T | RING | GACTGTGCCaccTGTCTGGAGAAG | GGACGTGGAGCTGCTGCT | 68°C |
| C296S | RING | TGTGCCATCtctCTGGAGAAGTACATTG | GTCGGACGTGGAGCTGCT | 66°C |
| E304D | RING | TTGATGGAgacGAGCTGCGGG | TGTACTTCTCCAGACAGATG | 62°C |
| E305K | RING | TGATGGAGAGaagCTGCGGGT | ATGTACTTCTCCAGACAGATGG | 64°C |
| R307Q | RING | GAGGAGCTGcagGTCATCCCC | TCCATCAATGTACTTCTCCAGAC | 65°C |
| R307W | RING | AGAGGAGCTGtggGTCATCCC | CCATCAATGTACTTCTCCAGACAG | 66°C |
| V308L | RING | GGAGCTGCGGctcATCCCCTG | TCTCCATCAATGTACTTCTCCAG | 64°C |
| P310S | RING | GCGGGTCATCtccTGTACTCA | AGCTCCTCTCCATCAATGTAC | 65°C |
| R314Q | RING | TGTACTCACcagTTTCACAGGAAGTGC | GGGGATGACCCGCAGCTC | 68°C |

| | | | | |
|---|---|---|---|---|
| H316Y | RING | TCACCGGTTTtacAGGAAGTGC | GTACAGGGGATGACCCGC | 66°C |
| R317M | RING | CGGTTTCACatgAAGTGCGTG | GTGAGTACAGGGGATGAC | 62°C |
| V320M | RING | CAGGAAGTGCatgGACCCCTG | TGAAACCGGTGAGTACAGG | 65°C |
| D321N | RING | GAAGTGCGTGaacCCCTGGCT | CTGTGAAACCGGTGAGTACAG | 66°C |
| P322S | RING | GTGCGTGGACtccTGGCTGCT | TTCCTGTGAAACCGGTGAGTACAGG | 72°C |
| C330Y | RING | CACCACACCtacCCCCACTGTC | CTGCAGCAGCCAGGGGTC | 70°C |
| P331H | RING | CACACCTGCcacCACTGTCGG | GTGCTGCAGCAGCCAGGG | 72°C |
| P331L | RING | CACACCTGCctcCACTGTCGG | GTGCTGCAGCAGCCAGGG | 72°C |
| C333R | RING | CTGCCCCCACcgtCGGCACAA | GTGTGGTGCTGCAGCAGC | 71°C |
| C333S | RING | CTGCCCCCACagtCGGCACAA | GTGTGGTGCTGCAGCAGC | 71°C |
| V228M | TM | TTTCTTCGTCatgGTCTCCTTG | GCCAGGAAAATCCCCATG | 61°C |
| S230F | TM | GTCGTGGTCttcTTGGTCTGC | GAAGAAAGCCAGGAAAATCCC | 64°C |
| P369L | DIR | GTGCATTACctcGGCCGCGTG | CGGCAGGGTCACCCTCTG | 72°C |
| P382S | DIR | CCCAGCCTACtctACGAGGAC | ATGGCGTTGGTCCTGTGC | 69°C |
| H533N | other | CAGCTGCTATaacGGCCACCG | AAGCTGGACGTGCTGCCG | 71°C |
| A544V | S-rich | GGCTACCTGgtcGACTGCCCA | ACTGCACACCGAGCGGTG | 72°C |
| D545N | S-rich | CTACCTGGCCaacTGCCCAGG | CCACTGCACACCGAGCGG | 71°C |
| S683L | other | AGCTCTACCttaGAAGTGGGGC | ATTGGGCCCCCTGTGCTC | 67°C |

### Supplementary Table 4. Primers used to introduce missense mutations in the *RNF43* gene

| Mutation | Domain | Forward primer | Reverse primer | Tm |
|---|---|---|---|---|
| C275F | RING | TGTGCCATCTtTCTGGAGGAGTTCTC | CACAGGGGCTGAGCTGCA | 71°C |
| L276P | RING | GCCATCTGTCcGGAGGAGTTCTC | ACACACAGGGGCTGAGCT | 71°C |
| E277K | RING | CATCTGTCTGaAGGAGTTCTCTGAGG | GCACACACAGGGGCTGAG | 69°C |
| F279L | RING | TCTGGAGGAGcTCTCTGAGGG | CAGATGGCACACACAGGG | 67°C |
| E281K | RING | GGAGTTCTCTaAGGGGCAGGA | TCCAGACAGATGGCACAC | 66°C |
| Q283R | RING | TCTGAGGGGCgGGAGCTACGG | GAACTCCTCCAGACAGATGGCAC | 71°C |
| L285I | RING | GGCAGGAGaTACGGGTCATTTC | CCTCAGAGAACTCCTCCAGACAGATG | 69°C |
| R286L | RING | CAGGAGCTACtGGTCATTTCC | CCCCTCAGAGAACTCCTC | 65°C |
| V287F | RING | GGAGCTACGGtTCATTTCCTGC | TGCCCCTCAGAGAACTCC | 68°C |
| V287A | RING | GAGCTACGGGcCATTTCCTGC | CTGCCCCTCAGAGAACTC | 66°C |
| S289F | RING | CGGGTCATTTtCTGCCTCCATG | TAGCTCCTGCCCCTCAGA | 69°C |
| C290Y | RING | GTCATTTCCTaCCTCCATGAGTTCCATCG | CCGTAGCTCCTGCCCCTC | 71°C |
| H292Y | RING | TTCCTGCCTCtATGAGTTCCATC | ATGACCCGTAGCTCCTGC | 67°C |
| E293G | RING | TGCCTCCATGgGTTCCATCGT | GGAAATGACCCGTAGCTC | 64°C |
| E293D | RING | GCCTCCATGAtTTCCATCGTAAC | AGGAAATGACCCGTAGCT | 64°C |
| F294S | RING | CTCCATGAGTcCCATCGTAAC | GCAGGAAATGACCCGTAG | 64°C |
| C298F | RING | CATCGTAACTtTGTGGACCCC | GAACTCATGGAGGCAGGA | 65°C |
| V299M | RING | TCGTAACTGTaTGGACCCCTG | TGGAACTCATGGAGGCAG | 65°C |

| | | | | |
|---|---|---|---|---|
| D300A | RING | AACTGTGTGGcCCCCTGGTTA | ACGATGGAACTCATGGAG | 62°C |
| D300V | RING | AACTGTGTGGtCCCCTGGTTA | ACGATGGAACTCATGGAG | 62°C |
| W302R | RING | TGTGGACCCCaGGTTACATCA | CAGTTACGATGGAACTCATG | 61°C |
| Q305K | RING | CTGGTTACATaAGCATCGGAC | GGGTCCACACAGTTACGA | 64°C |
| R307Q | RING | CATCAGCATCaGACTTGCCCC | TAACCAGGGGTCCACACA | 67°C |
| C309Y | RING | CATCGGACTTaCCCCCTCTGC | CTGATGTAACCAGGGGTC | 63°C |
| C309W | RING | ATCGGACTTGgCCCCTCTGCA | GCTGATGTAACCAGGGGTCC | 69°C |
| P310H | RING | CGGACTTGCCaCCTCTGCATG | ATGCTGATGTAACCAGGGGTC | 68°C |
| L311C | RING | GACTTGCCCCaTCTGCATGTT | CGATGCTGATGTAACCAGG | 64°C |

### Supplementary Table5. Primers used for qPCR

The following primers were used to compare the different ZNRF3 isoforms with qPCR:

| **ZNRF3 Isoform** | **Primer type** | **Sequence 5' →3'** |
|---|---|---|
| Short | Pair 1, forward | GGAGTGTCCTGTGTGGTGTCC |
| Short | Pair 1, reverse | GGCCTCTTGAGCGGGTCTTC |
| Short | Pair 2, forward | TGTCCTGTGTGGTGTCCACTT |
| Short | Pair 2, reverse | GCTTCTGGGTTTTCAGACACATC |
| Long | Pair 1, forward | AGGGCGAGATCGTGCAGATG |
| Long | Pair 1, reverse | GGCCTCTTGAGCGGGTCTTC |
| Long | Pair 2, forward | GGCGAGATCGTGCAGATGC |
| Long | Pair 2, reverse | GCTTCTGGGTTTTCAGACACATC |

As controls, the following primer pairs specific for both ZNRF3 isoforms (Ex8/9) and specific for the housekeeping gene GAPDH were also included:

| **Primer** | **Sequence 5' → 3'** |
|---|---|
| ZNRF F | TGCTGTCAGGGCCAACTT |
| ZNRF R | CAGTTCCCAATTTCCAGGTAAG |
| GAPDH m103 F | AGAGGGATGCTGCCCTTACC |
| GAPDH m103 R | GTAAAGCCGCGAGTAGCTGG |

The following primers were used to compare the overexpression of ZNRF3 and RNF43 with qPCR:

| **Primer** | **Sequence 5' → 3'** |
|---|---|
| ZNRF F1 | GGGTCATCCCCTGTACTCAC |
| ZNRF R1 | TTGTCCTCGTAGGGTAGGCTG |
| RNF43 F | ATCAGCATCGGACTTGCCC |
| RNF43 R | GGATGCTGGCGAATGAGGTG |

### Supplementary Table6. gRNAs used for CRISPR/Cas9 genome editing and the primers used for repaired templates

| ZNRF3 mutations | gRNA | Sequence 5' →3' |
|---|---|---|
| P179L | forward | CACC GCACCCTTCACATACACCAC |
| | reverse | AAAC GTGGTGTATGTGAAGGGTGC |
| R307W | forward | CACC gAGGGGATGACCCGCAGCTCC |
| | reverse | AAAC GGAGCTGCGGGTCATCCCCTc |
| P331L/H & C333R | forward | CACC gACCACACCTGCCCCCACTGT |
| | reverse | AAAC ACAGTGGGGGCAGGTGTGGTc |
| R246* | forward | CACC GCTGAAGCAGCGACGCAGTC |
| | reverse | AAAC GACTGCGTCGCTGCTTAAGC |
| S355* | forward | CACC GAGACCAGCAACCTCTCACG |
| | reverse | AAAC CGTGAGAGGTTGCTGGTCTC |
| R536* | forward | CACC GAGCCACTGCACACCGAGCGG |
| | reverse | AAAC CCGCTCGGTGTGCAGTGGCTC |
| R789* | forward | CACC GAGAAGCAGGTGGCCCGCGG |
| | reverse | AAAC CCGCGGGCCACCTGCTTCTC |

| ZNRF3 mutations | primers used for repaired templates | Sequence 5' →3' |
|---|---|---|
| P179L | forward | AGTATATGTGAAGGGTGCAGATGC |
| | reverse -WT | ACCGGCCTCTTGAGCGGGTCTT |
| | reverse -MUT | ACCAGCCTCTTGAGCGGGTCTT |
| R307W | forward-WT | ACGGGTCATCCCCTGTACTCACCG |
| | forward -MUT | ATGGGTCATCCCCTGTACTCACCG |
| | reverse | AGTTCCTGGGCAGGGAGAGAGGG |
| P331L/H | forward | GCCGTCACAACATCATAGGTAACTGTCACCCG |
| | reverse -WT | AGTGGGGGCAGGTGTGGTGCTGCA |
| | P331L-reverse -MUT | AGTGGAGGCAGGTGTGGTGCTGCA |
| | P331H-reverse -MUT | AGTGGTGGCAGGTGTGGTGCTGCA |
| C333R | forward | GTCGTCACAACATCATAGGTAACTGTCACCCG |
| | reverse -WT | AGTGAGGGCAGGTGTGGTGCTGCA |
| | reverse -MUT | GGTGAGGGCAGGTGTGGTGCTGCA |

### Supplementary Table7. Primers used for identifying genome-edited mutants generated with CRISPR/Cas9

The following primers were used to screen for sigle cells clones that cotained the desired truncating mutations. Primers names containing 'in' refer to the innner primers. Noprimeris given for R789 truncation as this mutation was screened using a Cfr42I(alias SacII) digestion.

| Primer | Sequence 5' → 3' |
|---|---|
| ZNRF3-R246-scF | GGAAGGGCCTTTCTTACCAGT |

| | |
|---|---|
| ZNRF3-R246-scR | TGCTGGAGATATTCCCAATGCTT |
| ZNRF3-R246-scFin | TGAAGCAGCGACGCAGT |
| ZNRF3-Ex8F | AATGGGTACCTTGGCAGGTG |
| ZNRF3-1397R | TCATACTGGGAGAAGCAAGCTG |
| ZNRF3-S355-scFin | GGAGACCAGCAACCTCTCAC |
| ZNRF3-S536-scF2 | CCCTACGAGGACAAGCATGG |
| ZNRF3-S536-scRin2 | TGCTATCACGGCCACCGCT |
| ZNRF3-S536-scR2 | GGAGGAGTTGCTGCTGTAGT |
| ZNRF3-G789-scF | CCCTCAGCAGCGACTATGACC |
| ZNRF3-G789-scR | AGGGCACTAGGGAAGTTGGC |
| ZNRF3-gDNA-ex7F1 | ATGAGCAAATCTGGAAGATG |
| P179L-MUT-F | AAGACCCGCTCAAGAGGCt |
| ZNRF3-gDNA-ex7R | TCGTGCAGACAACCTCCAAA |
| ZNRF3-gDNA-ex7F | GGAGGTGGTTCTGTCCCAAT |
| R307W-MUT-scR2 | TGAGTACAGGGGATGACCCAT |
| ZNRF3-P382S-R | ATGGCGTTGGTCCTGTGC |
| ZNRF3-gDNA-ex7R | TCGTGCAGACAACCTCCAAA |
